# Time-resolved directional organization of brain networks: distinct strength and temporal signatures from neurophysiology to disease

**DOI:** 10.64898/2026.02.15.706049

**Authors:** Nan Xu, Xiaodi Zhang, Wen-Ju Pan, Jeremy L. Smith, Eric H. Schumacher, Jason W. Allen, Vince D. Calhoun, Shella D. Keilholz

**Affiliations:** Fischell Department of Bioengineering, Department of Electrical and Computer Engineering, Brain and Behavior Institute, University of Maryland, College Park, MD, United States; Wallace H. Coulter Department of Biomedical Engineering, Georgia Institute of Technology and Emory University, Atlanta, GA, United States; Department of Radiology and Imaging Sciences, Emory University School of Medicine, Atlanta, GA, United States; School of Psychology, Georgia Institute of Technology, Atlanta, GA, United States; Department of Radiology and Imaging Sciences, Indiana University School of Medicine, Indianapolis, IN, United States; Tri-institutional Center for Translational Research in Neuroimaging and Data Science, Georgia State University and Georgia Institute of Technology and Emory University, Atlanta, GA, United States

## Abstract

Brain function depends on time-resolved directional organization: influence is continuously redistributed across regions, shaping who drives whom, when, and for how long. Yet dynamic neuroimaging has largely emphasized undirected fluctuations in functional connectivity, limiting biological interpretation of asymmetric interactions. Here we show that directed influence can be resolved into two biologically distinct signatures: strength, reflecting the efficacy of influence, and duration, reflecting its temporal extent. To test this principle, we developed sliding-window prediction correlation (SWpC), a framework for estimating these signatures in fMRI and related signals. Across simultaneous rat local field potential (LFP)-fMRI and human motor-task fMRI, strength more closely tracks neural-BOLD coupling and task-evoked reconfiguration, whereas duration captures temporally extended BOLD patterns more susceptible to hemodynamic timing. In post-concussion vestibular dysfunction, altered strength-duration profiles define reproducible vestibular-multisensory states and improve patient-control discrimination. Together, these findings support a general principle of dynamic brain organization spanning neurophysiology, behavior, and disease.

## 1. Introduction

The brain is a dynamic system in which interactions among regions are continually reconfigured over time (Allen et al., 2012; R M Hutchison et al., 2013). But brain networks are not only time-varying; they are also directionally organized (Friston, 2011; A Mitra et al., 2015; Xu et al., 2021b). At any moment, function depends not just on whether regions are coupled, but on who influences whom, when that influence emerges, and how long it persists. This time-resolved directional organization is central to biological computation because it supports selective routing, temporal coordination across distributed systems, and rapid redistribution of influence as behavioral demands change. Understanding brain function therefore requires tracking not only changing coupling, but changing directed influence across neurophysiology, behavior, and disease.

Despite this, most dynamic neuroimaging studies still emphasize undirected time-varying functional connectivity. Sliding-window correlation (SWC) and related approaches have been valuable for identifying recurring network states and linking them to cognition and disease, but they remain fundamentally symmetric: they capture whether signals covary within a window, not whether one region preferentially influences another (Allen et al., 2012; Shakil et al., 2016). This limits mechanistic interpretability, especially when the biological question concerns asymmetric interactions, hierarchical organization, or altered information routing in disease, including conditions such as post-concussion vestibular dysfunction (PCVD) (Smith et al., 2021). As a result, an important feature of dynamic brain organization—time-resolved directionality—remains under-characterized in widely used neuroimaging frameworks.

A key reason directionality matters biologically is that “directed influence” is not a single quantity. Neural interactions can differ not only in how strongly one system drives another, but also in how long that influence persists. Yet many lag-based directed methods assume a fixed lag/history structure, effectively treating interaction timescales as constant across time and condition. This obscures the elasticity of brain interactions, i.e., how the effective lag span expands or contracts with state, and motivates window- and direction-specific duration estimates learned from the data (Park et al., 2026; Seth et al., 2015; Stephen M. Smith et al., 2011). A brief, high-gain interaction can reflect rapid, event-like control or gating, whereas a weaker but temporally sustained influence can reflect integration, recurrent reverberation, or propagation along slower pathways (Chaudhuri et al., 2015; Fries, 2015; Anish Mitra et al., 2015). These two aspects—strength and temporal persistence—are expected to have distinct biological determinants and may dissociate across contexts (Murray et al., 2014; Xu et al., 2021a). Critically, they may also be shaped differently in fMRI: hemodynamic filtering can smear and delay signals, potentially altering the apparent temporal footprint of influence even when underlying neural coupling is unchanged (Handwerker et al., 2004; R. Matthew Hutchison et al., 2013). This motivates a biology-forward framework that can resolve directed interactions into separable signatures of strength and temporal duration.

Historically, neuroimaging methods for correlation and causality have been treated as conceptually and practically distinct (Aedo-Jury et al., 2020; Friston, 2011; Friston et al., 2003; Keilholz et al., 2013; Liang et al., 2015; Park et al., 2018; van den Heuvel and Hulshoff Pol, 2010). Correlational approaches quantify statistical dependence without directionality, whereas causal models aim to identify asymmetric influence by distinguishing drivers from recipients (Peters et al., 2017). Recent advances have enabled time-varying effective connectivity in EEG/MEG, often in low- to moderate-dimensional settings, e.g., (Medrano et al., 2024). Complementary developments are still needed to enable scalable, whole-brain, time-resolved modeling for resting-state fMRI, where hemodynamic filtering and high dimensionality impose distinct constraints. This practical gap has constrained efforts to integrate correlational and directed perspectives within a unified, time-resolved framework.

To address this practical gap, we developed sliding-window prediction correlation (SWpC), a computational framework that retains the practical sliding-window workflow while enabling time-resolved directionality and the resolution of directed influence into distinct signatures of strength and duration. Within each window, SWpC embeds an impulse-response prediction model to predict a target timeseries from a source in a specified direction (x→y). This windowed directed modeling resolves time-resolved directed influence into two biologically distinct signatures: directed strength, defined as the predictive efficacy of x→y within a window, and directed duration, defined as the window-specific temporal lag footprint over which that influence persists. Importantly, strength and duration need not covary—interactions can be brief but strong, or weaker yet temporally sustained—supporting a biologically motivated decomposition that remains scalable for whole-brain fMRI analyses.

Because directionality inference from BOLD fMRI can be confounded by regional variability in hemodynamic responses, biological validation is essential. Simulation studies are useful for probing assumptions (S M Smith et al., 2011; Xu et al., 2017), but they cannot fully capture the constraints of in vivo neurophysiology and neurovascular coupling. We therefore evaluate SWpC across complementary benchmarks spanning multiple levels: simultaneous rodent LFP–fMRI to link directed signatures in neuronal activity and BOLD, and human motor-task fMRI to test sensitivity to behaviorally driven reconfiguration of directional organization. These datasets test a central biological hypothesis: that time-resolved directional organization is expressed through two distinct signatures, i.e., strength and duration, which show convergent structure across neurophysiology and behavior while being differentially shaped by hemodynamic filtering.

Finally, we use post-concussion vestibular dysfunction (PCVD) as a clinical demonstration of translational utility. Vestibular symptoms reflect distributed interactions among vestibular, multisensory, and control systems, making PCVD a compelling setting in which disrupted directional organization may manifest as altered time-resolved network states (Smith et al., 2021). We test whether SWpC-derived signatures identify reproducible vestibular–multisensory brain states and improve patient–control discrimination relative to undirected dynamic FC, positioning the clinical cohort as a demonstration of practical relevance rather than the primary basis for mechanistic generalization.

Together, this work advances a biology-forward view of dynamic brain networks: time-resolved directional organization is a core principle of brain function, and it can be mechanistically interpreted by disentangling directed influence into strength and duration. SWpC provides a unified, scalable framework for estimating these signatures in fMRI, enabling analyses that connect neurophysiology, behavior, and neurological disease.

## 2. Results

### 2.1. Sliding-window prediction correlation (SWpC) framework for time-resolved directional connectivity

The model integrates the sliding-window technique with a validated static causal model of prediction correlation (Nan Xu et al., 2017; Xu et al., 2021) to analyze time-varying causal interactions in brain networks. Conceptually, in the causal and time-varying functional brain networks (FBNs), every directed functional connectivity (FC) interaction (e.g., x→y and y→x) is modeled as an independent communication channel within a causal, time-varying system.

To ensure computational scalability, the sliding-window correlation (SWC) approach approximates the time-varying system by dividing the time course into short, overlapping windows, each treated as a time-invariant model. Within these windows, a finite impulse-response (FIR) model (e.g., see framework as outlined in (Nan Xu et al., 2017; Xu et al., 2021)) is embedded for analysis. For a specific window [*τ*, *τ* + *L*] (starting at time τ with length L), the directed FC can be computed in the following two steps:

1. Prediction: The windowed segment of y (denoted by *y*_*τ*_) can be predicted by an FIR model, 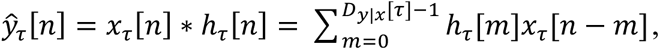, for *n* ∈ [*τ*, *τ* + *L*], where *x*_*τ*_ is an x segment within the current and the previous windows, and ℎ_*τ*_ [*n*] is the window-specific impulse response with duration *D_y_*_|*x*_ [*τ*]
2. Correlation: The strength of directed FC is calculated as the correlation between the predicted and observed segments: *ρ_y_*_|*x*_ [*τ*] = *corr*(*ŷ*_*τ*_, *y*_*τ*_).

Notably, in step (1), the input signal x in the previous window is also included for capturing the causal influences (see Fig. 1). Due to the previous success of prediction correlation in (Nan Xu et al., 2017; Xu et al., 2021), the FIR system can be reliably predicted for each window. The model predicts time-varying directed FC, with both strength (*ρ_y_*_|*x*_[*τ*]) and duration (*D_y_*_|*x*_[*τ*]) depending on the window’s starting point τ, enabling directional distinctions (*ρ_y_*_|*x*_[*τ*]≠*ρ_x_*_|*y*_[*τ*] and *D_y_*_|*x*_[*τ*]≠*D_x_*_|*y*_[*τ*] for any *τ* ’s). This dynamic method, namely sliding window prediction correlation (SWpC), extends SWC by incorporating causal influences and distinguishing between forward and backward connections, offering a more nuanced analysis of brain connectivity.

**Fig. 1.**
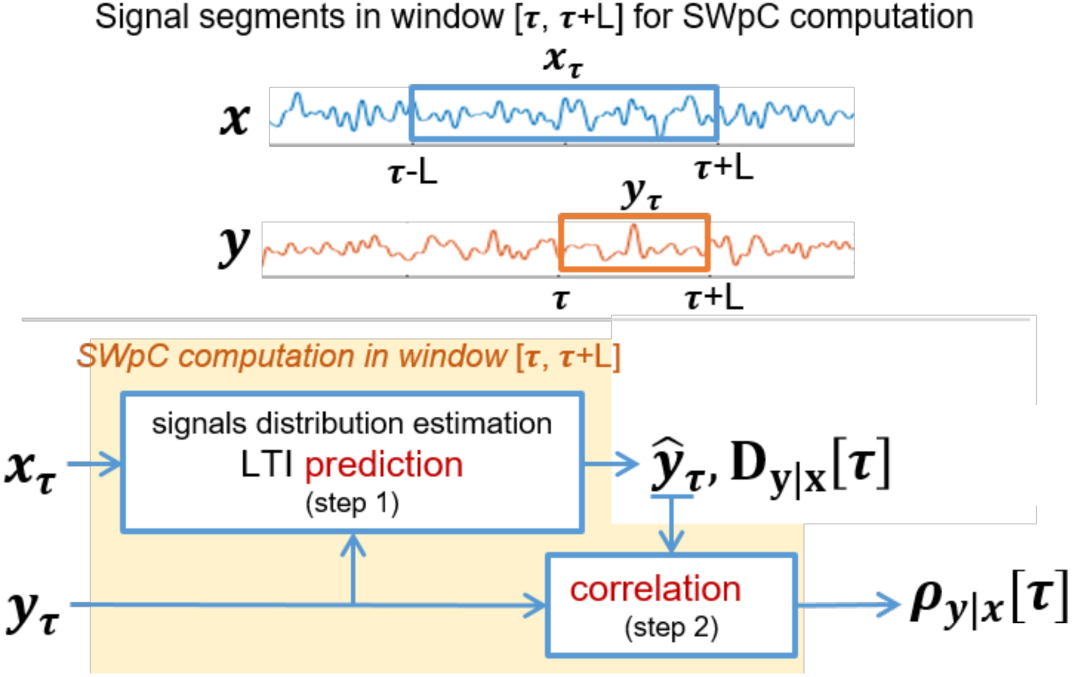
SWpC computation in each window.

### 2.2. Directed strength and duration show distinct biological grounding in simultaneous LFP–BOLD

The strength and duration of SWpC were reliably estimated from both BLP and BOLD signals recorded in S1L and S1R of rats (Fig. 2a), highlighting symmetrical connectivity between the two regions. Notably, the 75% quantile of directional asymmetry remained lower than the scan-to-scan variability (Fig. 2b). Figure 2B shows the distribution of mean asymmetry values across scans for each signal type, with the scan variability overlaid as a reference line. P-values derived from the Wilcoxon signed-rank test were all well below 0.05, confirming that the observed asymmetries in both strength and duration measures were significantly smaller than the scan variability, thus attesting to the stability of SWpC estimates. Furthermore, SWpC strength captured LFP-BOLD relationships similar to those revealed by SWC, showing strong BLP-BOLD correlations within the θ, low β, and higher frequency bands (with mean correlations >0.1, Fig. 2c). This finding is consistent with prior results obtained from concurrently recorded LFP-BOLD data in rats under isoflurane anesthesia (Garth John Thompson et al., 2013a). In contrast, SWpC duration displayed comparatively modest LFP-BOLD correlations (with mean correlations <0.1, Fig. 2c).

**Fig. 2.**
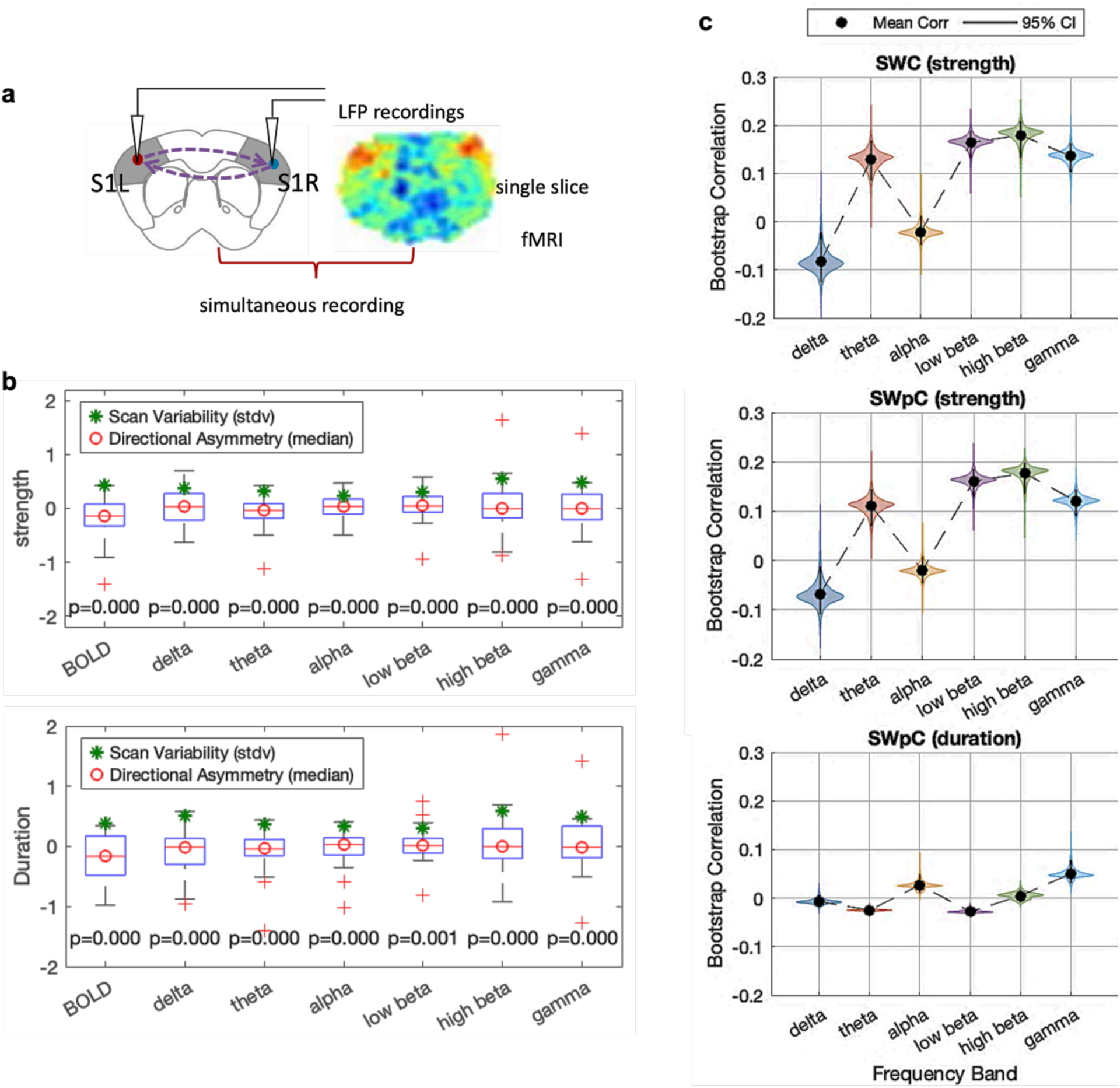
Reliability of the SWpC Measure in Simultaneously Recorded LFP-BOLD Data. **a** Schematic of simultaneous LFP and BOLD signal recordings from bilateral somatosensory regions (S1L and S1R). **b** Boxplots comparing directional asymmetry in connectivity (S1L→S1R vs. S1L←S1R) with scan-to-scan variability in SWpC strength (left) and SWpC duration (right) estimates for each signal type. The red line and red circle indicate the median directional asymmetry within each boxplot, while the green star denotes the corresponding scan-to-scan variability. **c** Bootstrap correlation analysis between BOLD results and band-limited power (BLP) results in six frequency bands for three metrics: SWC strength (left), SWpC strength (middle), and SWpC duration (right). For SWpC result, S1L→S1R is used as an illustrative example.

### 2.3. Task-evoked directional organization in human motor networks

As described in the Method section, the data were obtained from 45 subjects in the HCP test-retest dataset, where participants were cued to move their left or right foot, hand, or tongue during two repeated runs. Following the 3-level Feat analysis, we identified 49 ROIs (listed in Fig. S3) that align with previously reported motor task activation maps and connectome fingerprints covering the s (Barch et al., 2013; Tripathi et al., 2024). Specifically, foot activations are located on the superior midline, hand activations ventral to the foot, and tongue activations ventral to the hand (Fig. S2–4). In the cerebellum, ipsilateral hand/foot and bilateral tongue representations contrast with the contralateral patterns observed in the motor cortex.

As shown in Fig 3a and Fig S5, the SWpC results demonstrate that motion enhances directional information flow compared to the corresponding rest trials, which appear to exhibit more symmetrical connectivity patterns as shown in SWC results. This enhancement is evident in both strength and duration, with SWpC uniquely capturing temporal aspects of directed connectivity that are not addressed by SWC. Across all motion types, directional asymmetry in strength and duration during task performance was significantly greater than during rest, with Cohen’s *d* values exceeding 0.8 (see mean asymmetry and Cohen’s *d* values in Table S1), indicating large effect sizes and substantial differences between the two conditions. The violin plots in Fig. 3b further emphasize these findings, showing consistently increased asymmetry in directed FC during tasks relative to rest, in both strength and duration, across all motions.

**Fig. 3.**
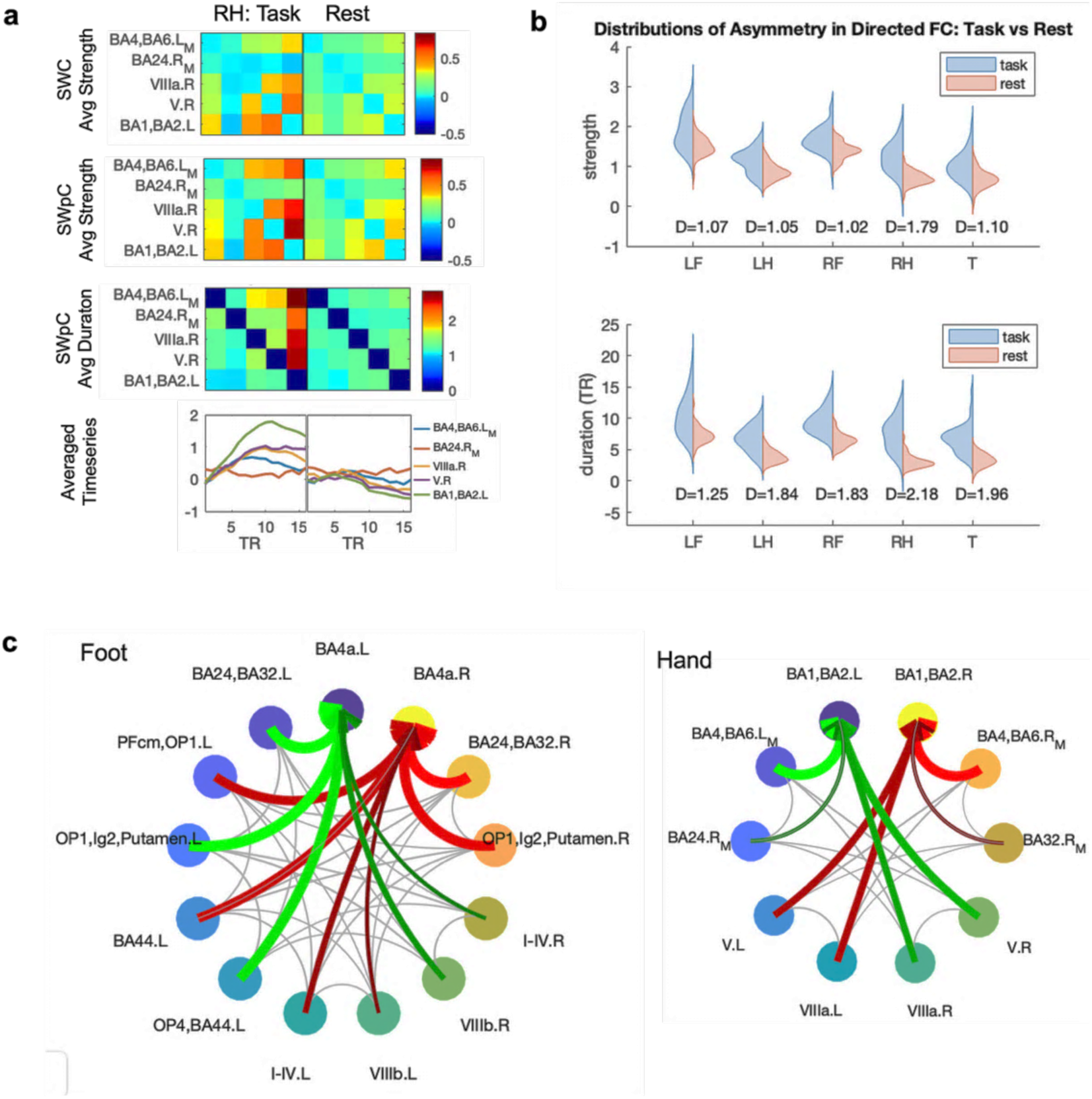
Asymmetry in directed FC and significant long durations between task vs. rest. **a** Example of group-averaged SWpC results for right-hand (RH) task, illustrating enhanced directional information flow compared to rest trials. All task-specific results are presented in Fig. S5. The top row presents the averaged FC strength from sliding window correlation (SWC). The second and third rows display the averaged directed FC strength and duration estimated by sliding window prediction correlation (SWpC), respectively. For simplicity, only larger right-hand ROIs (≥25 voxels) are included. The bottom row shows the averaged time series across all task (or all rest) trials for the five right-hand regions of interest (ROIs). **b** Violin plots summarizing directional asymmetry in strength and duration for all tasks, showing consistently increased asymmetry during task performance relative to rest, with large effect sizes (Cohen’s d > 0.8). The corresponding Cohen’s *d* value is listed under each pair of violin plots, indicating the magnitude of the difference in means of task and rest distributions relative to their pooled standard deviation (with values ≥0.8 generally considered large effects). **c** Connectograms of directed FC with significant differences in duration between task and rest for hand (left) and foot (right) motions. Red (green) arrows indicate task vs. rest differences for left (right) motion with low individual variability (CV < 36%). Gray connections represent all predicted (directed) connections based on SWpC. For clarity, the connectograms include 13 ROIs for foot and 10 ROIs for hand motions, excluding smaller ROIs (fewer than 100 voxels for foot motions and fewer than 30 voxels for hand motions). Remaining ROIs demonstrate approximate symmetry for left/right foot and hand tasks. Under these thresholds, tongue-evoked information flow among the 4 larger ROIs does not form hubs in the motor cortex as shown in Fig. S6.

Directed FC durations predicted by SWpC revealed significant differences between task and rest trials (FDR-corrected p < 0.01). Specifically, directed FC with significantly longer durations during tasks performances, coupled with low individual variability (CV < 36%), identified hub-like structures associated with hand and foot movements (Fig. 3c). Large ROIs in somatomotor areas responding to these movements exhibited increased long-range information transfer from other ROIs, as indicated by red and green arrows highlighting significant differences for left and right motions, respectively. Additionally, the averaged time series for each task, across all trials and subjects, revealed that these ROIs exhibited delayed and higher hemodynamic peaks compared to other ROIs (Fig. 3a, bottom row, and Fig. S5c). This temporal delay may account for the observed increase in the duration of information transfer.

Significant differences between the motion task and rest trials were also observed in the SWpC-predicted directed FC strengths. Figure 4 highlights the significantly enhanced directed FC strength evoked by each task, characterized by low individual variability (FDR-corrected *p* < 0.01, CV < 30%), contrasting with the symmetrical patterns estimated by SWC. For each of the hand and foot tasks, SWpC detected significantly evoked causal interactions between the cerebellum and the contralateral somatomotor regions, aligning with the SWC undirected FC results. Examples include VIIIb.L → BA4a.R for left foot motion, VIIIb.R → BA4a.L for right foot motion, VIIIa.L → BA1, BA2.R for left-hand motion, and VIIIa.R → BA1, BA2.L for right-hand motion. Additionally, SWpC demonstrated its sensitivity in detecting information flow from the OP1, Lg2, and putamen to BA4a for both left and right foot movements. Consistent with current understanding of sensorimotor control and neural connectivity, these connections were more lateral for hand than foot movements and most lateral for the tongue movement task (Fieblinger, 2021; Rizzolatti and Luppino, 2001).

**Fig. 4.**
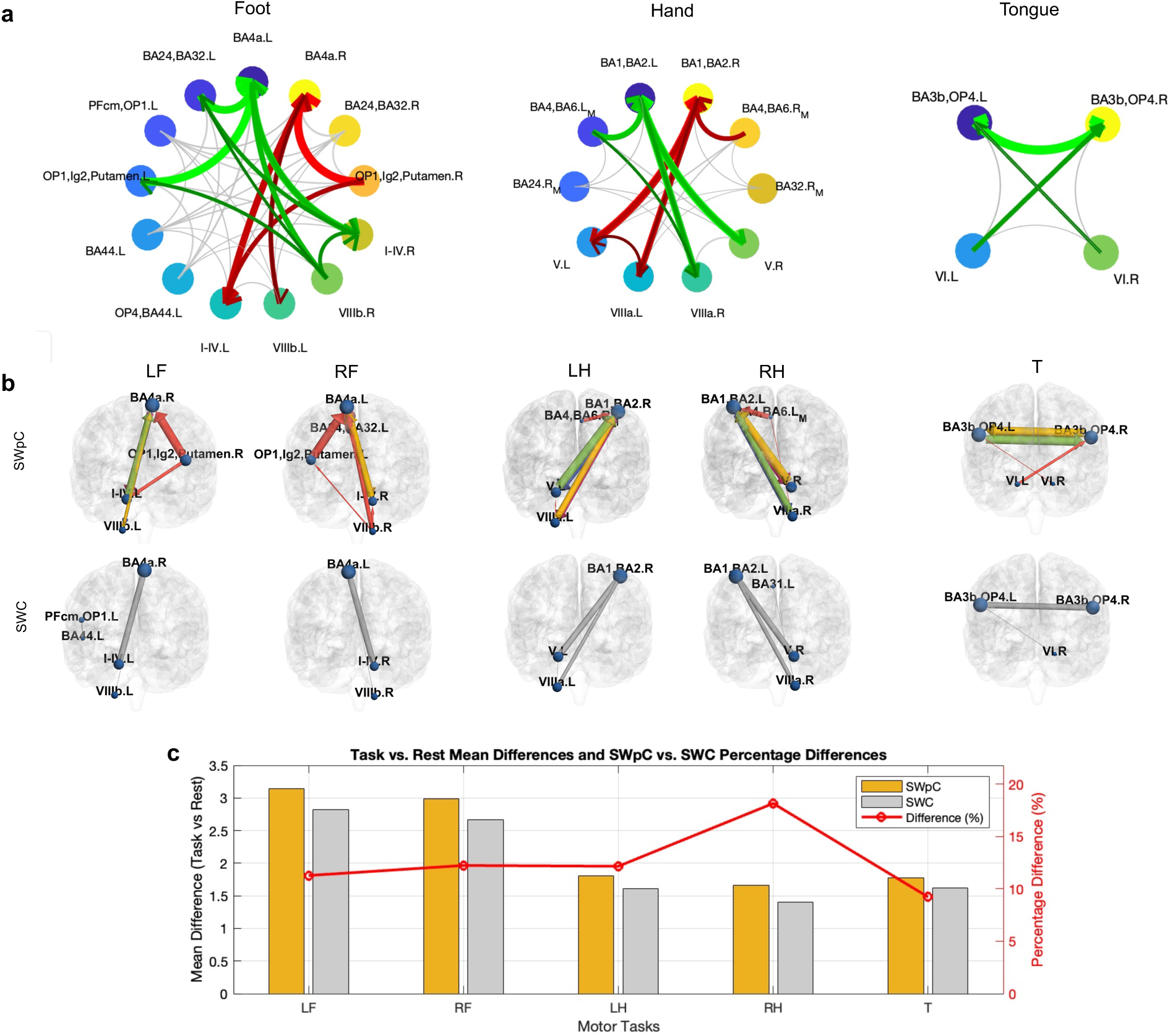
Directed FC with significant strength differences between task and rest for each motion. **a** Connectograms showing directed FC strengths significantly evoked by each motion (FDR-corrected p < 0.01). For hand and foot motions, red (green) arrows depict significant directed FC strengths for left (right) motions with low individual variability (CV < 30%). For tongue motions, green arrows represent significant evoked directed FC strengths with low variability. Gray connections denote all predicted directed connections from SWpC. To enhance clarity, connectograms include 13 foot ROIs, 10 hand ROIs, and 4 tongue ROIs, excluding smaller ROIs (fewer than 100 voxels for foot motions and fewer than 30 voxels for hand and tongue motions). The remaining ROIs exhibit approximate symmetry for left/right foot and hand tasks. **b** Brain regions and directed functional connections with significantly task-evoked strength predicted by SWpC are visualized and contrasted with SWC results. For both methods, connections with significant task-rest strength differences (FDR-corrected p < 0.01) and low variability (CV < 30%) are shown. For SWpC, these connections correspond to the red and green arrows in the connectogram in **a**. Arrow thickness represents connectivity strength, while the arrow color (cool to warm) indicates connection duration from short to long in SWpC results. **c** A bar plot comparing the mean strength differences between task and rest estimates for SWpC and SWC. The percentage difference, calculated as (SWpC−SWC)/SWC×100%, is shown as a red line.

The observed information flow from the medial extensional region of BA4/BA6 to BA1/BA2 during hand movements aligns with established evidence of the motor-sensory integration necessary for precise motor execution. This flow reflects the hierarchical motor control process, where motor planning and execution (BA4/BA6) are tightly coupled with sensory feedback processing (BA1/BA2) (Todorov, 2004). The ability of SWpC to detect such task-evoked directed functional connectivity underscores its sensitivity and reliability in identifying biologically meaningful information flow, as highlighted in effective connectivity studies of hierarchical motor control (Fieblinger, 2021).

Moreover, SWpC sensitively detected bilateral information flows from the cerebellum to contralateral tongue-specific regions in the somatomotor cortex (VI.L → BA3b, OP4.R; VI.R → BA3b, OP4.L), whereas SWC identified only one pair under the same p-value and CV thresholds. These findings are consistent with the well-established role of the cerebellum in coordinating fine motor task (Manto et al., 2011), including precise bilateral movements required for tongue control (Sasegbon and Hamdy, 2023). By capturing these subtle dynamics, SWpC demonstrates superior sensitivity, with over 10% greater detection of task-evoked connections compared to SWC (Fig. 4c).

Furthermore, temporal variations in SWpC strength were strongly correlated with SWC strength variations across all motion tasks (Pearson’s r = 0.85, p < 0.001), as shown in the scatter plot (Fig. S7). This strong correlation demonstrates consistency between the two methods in capturing FC strength temporal dynamics. The correlation between the group-averaged mean FC strength (calculated as the average of all connections within the group-averaged FC matrix) across the four trials (Fig. S8) and the group-averaged reaction time (RT) for each task is presented in Table 1. Despite the small sample size (n = 4) and the limited variability in RTs for these motions (Fig. S8), SWpC consistently demonstrated a reverse correlation with RT across all motion tasks, aligning with expectations. Notably, SWpC demonstrated greater sensitivity compared to SWC, particularly in tasks such as LH and T, where SWC yielded marginal positive correlation with RT. These findings underscore the enhanced capability of SWpC in detecting task-specific connectivity changes while maintaining agreement with traditional methods.

**Table 1:** Correlation (p-value) between the subject-averaged FC strength and the mean reaction time for each motion type across four trials.

|  | LF | RF | LH | RF | T |
| --- | --- | --- | --- | --- | --- |
| SWpC | -0.64 (0.36) | -0.96 (0.04) | -0.36 (0.64) | -0.82 (0.18) | -0.09 (0.91) |
| SWC | -0.86 (0.14) | -0.96 (0.04) | 0.13 (0.87) | -0.97 (0.03) | 0.001 (1.00) |

### 2.4. Dynamic directed brain states and group discrimination in PCVD

Using sliding-window connectivity features (SWC-strength, SWpC-strength, and SWpC-duration), we identified five reproducible dynamic brain states that capture distinct whole-network configurations across the vestibular–multisensory ROI set (Fig. 5a). Across both strength- and duration-based representations, the state structure showed two complementary (asymmetric) pairs of configurations with opposing patterns of within- versus between-network coupling. The remaining state played a more “baseline” role, but its expression depended on the metric: for SWpC-duration, state 1 was characterized by comparatively weaker and more diffuse connectivity, whereas for the strength-based representations, state 1 instead emphasized stronger within-network connections. Together, these shared states provide a common state space for summarizing time-varying vestibular–multisensory network organization in both healthy controls and PCVD patients.

**Fig. 5.**
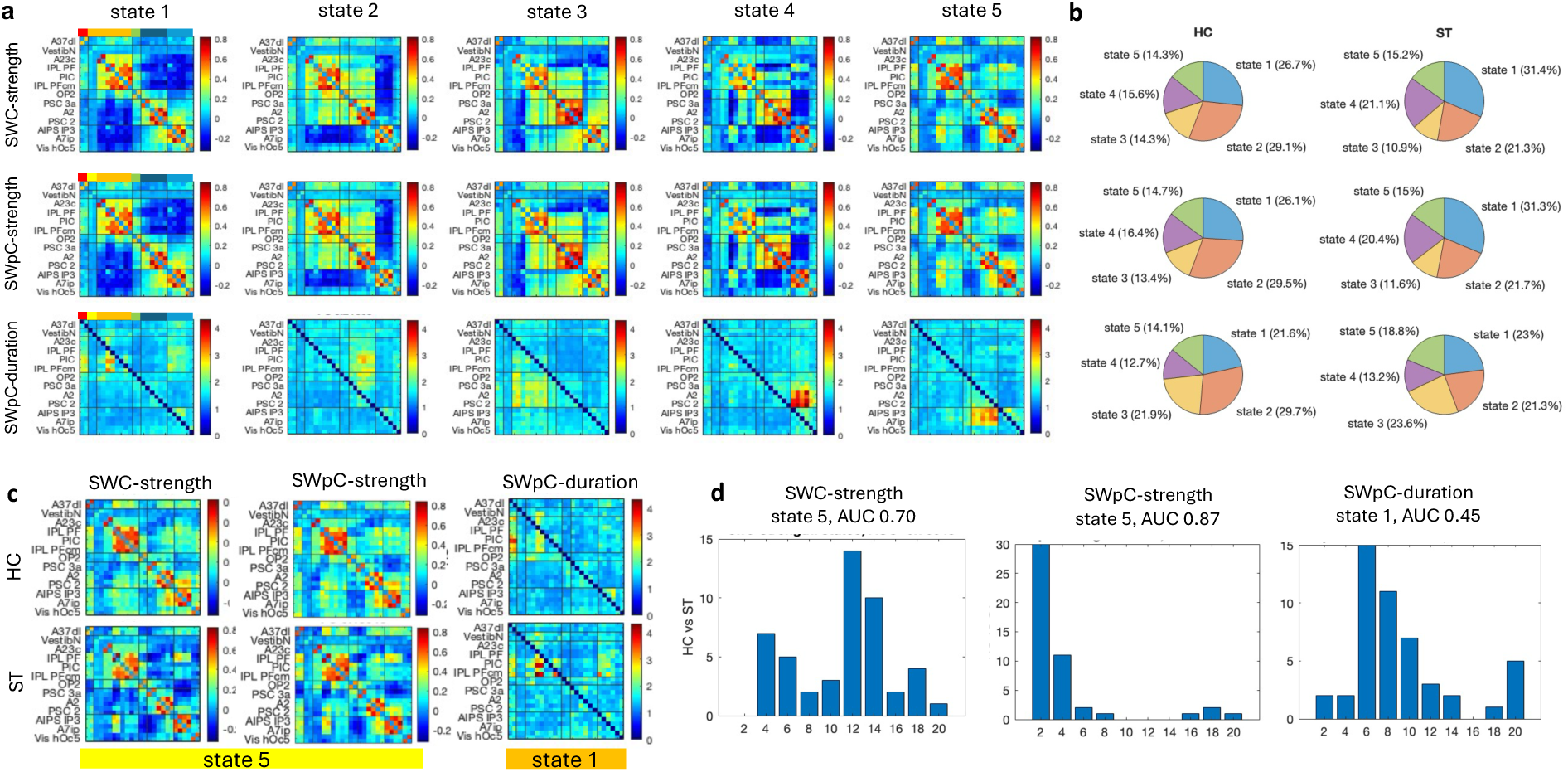
Brain-state dynamics and classification stability in PCVD. **a** Five recurring states derived from the full cohort (HC, ST, CH) for SWC-strength, SWpC-strength, and SWpC-duration. **b** Fractional occupancy in HC and ST for each measure. **c** Group-averaged matrices for the states with strongest HC–ST discrimination in nested LOOCV (state 5 for SWC-strength and SWpC-strength; state 1 for SWpC-duration). **d** Threshold-selection stability across nested LOOCV: bars show how often each threshold (thr = 1–20% in 2% steps) was selected; titles report the most frequently selected state and its AUC.

Notably, SWpC-strength resolved six communities within the core vestibular ROI set, whereas the traditional SWC-strength yielded only five communities and grouped the vestibular node (VestibN) with A23c, IPLPF, PIC, IPL, and PFcm (also see Fig. S12 for group specific brain states and networks). Given the vestibular node’s expected functional specificity, the finer community separation revealed by the SWpC-based measures appears more neurobiologically plausible and suggests that SWpC better preserves distinct vestibular-network organization (Frank and Greenlee, 2018). Fractional occupancy (FO) did not differ significantly between healthy controls (HC) and subacute PCVD (ST) for any state in SWC-strength (all p ≥ 0.12) or SWpC-strength (all p ≥ 0.13). In contrast, SWpC-duration showed a significant HC–ST difference in state 2 (p = 0.03), indicating that duration of causal interactions may capture complementary aspects of disease-related dynamics.

In nested LOOCV, the states that most strongly discriminated subacute PCVD (ST) from healthy controls (HC) were driven primarily by connectivity-strength features rather than duration. Specifically, state 5 was most frequently selected for both SWC-strength and SWpC-strength, yielding AUC ≈ 0.70 and AUC ≈ 0.87, respectively, based on the corresponding group-averaged state matrices (Fig. 5C). Notably, SWpC-strength was more sensitive than SWC-strength, as evidenced by its higher AUC for HC–ST discrimination. This pattern is consistent with the motion-task results in the last section, where SWpC-strength likewise demonstrated greater sensitivity than SWC-strength for detecting motion-evoked connectivity changes. In contrast, for SWpC-duration, the most frequently selected state 1 exhibited near-chance performance (AUC ≈ 0.45), suggesting limited discriminative value of duration-based features in this comparison (Fig. 5D).

## 3. Discussion

Time-resolved directional organization is a core feature of brain networks, linking neurophysiology to behavior and disease (R. Matthew Hutchison et al., 2013; Medrano et al., 2024; Xu et al., 2021a). Here we show that this organization can be interpreted through two complementary signatures of directed influence, i.e., strength and duration, that dissociate and are differentially shaped when estimated from BOLD fMRI (Anish Mitra et al., 2015; Xu et al., 2021a). We operationalize these signatures using sliding-window prediction correlation (SWpC), a scalable framework that estimates time-resolved directionality within the sliding-window workflow and yields directed strength (prediction correlation) and directed duration (model-selected impulse-response support). Across concurrent rodent LFP–BOLD recordings, human motor-task fMRI, and resting-state fMRI in PCVD, the convergent results support a general mechanism in which dynamic directional organization reconfigures with state, while the temporal extent of influence is more susceptible to hemodynamic timing effects than coupling strength. Taken together, the three datasets provide convergent evidence—from neurophysiology (LFP–BOLD) to behavior (motor task) to clinical dysfunction (PCVD)—that time-varying directed organization is a reproducible signature of brain network dynamics.

### 3.1. Reliability and biological grounding from simultaneous LFP–BOLD recordings

A key challenge for directed connectivity methods is establishing biological plausibility under realistic measurement conditions, particularly for BOLD signals that are shaped by hemodynamics. Using simultaneous LFP–BOLD recordings from bilateral somatosensory cortices, we validated SWpC against a strong physiological expectation: near-symmetric interhemispheric interactions between homologous regions (Moon et al., 2025). Both SWpC strength and duration satisfied this benchmark, with directional asymmetry consistently remaining below scan-to-scan variability. This indicates that SWpC does not spuriously impose directionality beyond what is supported by the data, and that its window-wise estimation is stable under realistic noise and nonstationarity.

Beyond stability, the multimodal recordings helped differentiate what SWpC strength and duration may capture when estimated from BOLD. SWpC strength showed BLP–BOLD correspondence consistent with prior SWC-based observations (stronger alignment in θ, low β, and higher-frequency bands), whereas SWpC duration exhibited comparatively weaker correspondence with band-limited power. A cautious interpretation is that BOLD-based duration is more susceptible to factors that shape the temporal profile of the fMRI signal such as hemodynamic smoothing, regional response timing differences, whereas BOLD-based strength more reliably reflects neural related covariation measured by LFP. This dissociation motivates treating duration as a complementary descriptor of temporal response structure rather than a direct proxy for “how long neural influence lasts.”

#### 3.2. Sensitivity to task-evoked directed interactions in human motor fMRI

In the HCP motor task, SWpC consistently revealed greater directional asymmetry during task than rest, indicating that task engagement produces more directionally organized interactions than those observed during the null condition. Importantly, SWpC detected task effects not only in directed strength but also in directed duration—an aspect that SWC cannot represent. Longer estimated durations formed hub-like patterns during hand and foot movements, suggesting that task engagement can expand the temporal footprint of directed interactions in addition to increasing their magnitude. Crucially, these detected information transfers were consistent with canonical sensorimotor circuit organization (e.g., cerebellar→somatomotor and effector-dependent lateralization) (Manto et al., 2011), supporting the neurobiological plausibility of SWpC-derived directionality.

At the same time, the duration findings are best understood as the BOLD timing. The longer durations likely reflect task-evoked hemodynamic timing effects: the time-averaged task-evoked BOLD responses in the task ROIs rise later and peak higher than those of other ROIs, consistent with a more temporally extended response profile (e.g., Fig. 3a). Hence, increasing SWpC duration even without implying a one-to-one mapping to prolonged neuronal influence. This interpretation aligns with the concurrent rat recordings, where duration showed weaker LFP–BOLD correspondence than strength. Taken together, these results suggest that, in task fMRI, SWpC duration is informative as a marker of temporally extended directed relationships in BOLD, potentially reflecting delayed or broadened responses in task-recruited regions, while SWpC strength may provide a more direct and stable index of directed coupling expressed in the signal.

SWpC strength additionally recovered task-evoked directed interactions that are coherent with known motor system organization, including cerebellar-to-contralateral somatomotor pathways and expected lateralization patterns across effectors. Under matched significance and variability thresholds, SWpC detected more task-evoked connections than SWC, supporting the view that prediction-based directionality can improve sensitivity while remaining neurobiologically plausible. Notably, SWpC and SWC strength variations remained strongly correlated across trials, indicating that SWpC preserves the core temporal dynamics captured by correlation-based approaches while adding interpretability in direction and temporal extent.

Finally, the exploratory association between group-averaged directed strength and reaction time consistently negative across tasks for SWpC, suggesting that stronger directed coupling may accompany faster performance. Given the small number of trial-level points (n=4), this analysis should be viewed as hypothesis-generating; nevertheless, the contrast with SWC (including marginal positive correlations in some tasks) motivates future studies with richer trial structure to test whether direction-aware measures better track behaviorally relevant network dynamics.

### 3.3. Dynamic vestibular–multisensory states in PCVD and implications for clinical utility

Applying SWpC to resting-state fMRI in PCVD patients, we identified a shared set of five reproducible vestibular–multisensory brain states across HC, subacute PCVD, and chronic PCVD (the latter included to stabilize state estimation). The state organization showed structured and interpretable configurations, including complementary asymmetric pairs that trade off within-versus between-network coupling, suggesting that directed dynamics provide a meaningful state space for summarizing vestibular–multisensory network organization.

Two findings are particularly relevant for clinical interpretation. First, SWpC-strength produced a finer modular decomposition within the vestibular ROI set than SWC-strength, separating the vestibular node into a distinct community rather than merging it with multiple parietal/insular parcels. Given the vestibular node’s expected functional specificity, this separation appears more consistent with known vestibular network organization (Frank and Greenlee, 2018; Indovina et al., 2020) and suggests that SWpC may better preserve functionally distinct pathways that are blurred in purely correlational representations.

Second, group differences depended on the feature type. Fractional occupancy did not differ significantly between HC and subacute PCVD for strength-based states, whereas duration-based measures showed a significant difference in one state. However, when assessing discriminability at the subject level, the strongest HC–ST separation emerged from strength-based features: state 5 was repeatedly selected as most discriminative for both SWC- and SWpC-strength, but SWpC-strength achieved substantially higher AUC than SWC-strength. In contrast, duration-based classification performance was near chance, which may reflect the greater variability of window-wise duration estimates and limited statistical power in this modest sample (n = 24 per group). Taken together, these results suggest that, in this dataset, directed strength carries the dominant discriminative signal, while duration may capture complementary group effects (e.g., occupancy shifts) without providing robust separability for classification. This pattern mirrors the motor-task results, where SWpC-strength also showed greater sensitivity than SWC-strength for detecting task-evoked changes, supporting the broader conclusion that prediction-based directed strength is a particularly informative and stable marker across contexts.

### 3.4. Methodological considerations and limitations

Several limitations should be considered when interpreting these findings. First, SWpC embeds a windowed impulse-response prediction model. While this choice supports interpretability and computational tractability, it is ultimately a proxy for directed interactions that may be nonstationary and/or nonlinear, with time-varying latency, state-dependent gain, or interaction shapes that are not well captured by a windowed impulse-response approximation. Relatedly, enforcing a fixed window length and overlap can constrain how rapidly the model adapts when interaction strength and temporal profile change at different rates across the scan. A promising direction is to incorporate adaptive window length so that the analysis can allocate shorter windows to rapidly changing regimes and longer windows to more stable periods. In parallel, extending SWpC to nonlinear time-varying causal system models (e.g., locally nonlinear predictors, kernelized or state-space formulations) could better capture elastic, context-dependent directed interactions while retaining the windowed workflow. Finally, the PCVD dataset is relatively small; although nested cross-validation mitigates information leakage, effect sizes and discriminability should be re-evaluated in larger cohorts.

### 3.5. Outlook

Overall, SWpC provides a practical expansion of the widely used sliding-window workflow by adding directionality and temporal extent of influence. Across multimodal validation and human task and patient applications, SWpC yields stable estimates, improves sensitivity to condition-dependent changes, and offers an interpretable decomposition of dynamic network organization. The observed dissociation between strength and duration further motivates treating these features as complementary axes of directed connectivity, which may prove useful for probing mechanistic hypotheses and for developing clinically relevant biomarkers as datasets grow at scale.

## 4. Methods

### 4.1. Simultaneously recorded LFP-BOLD data in lightly anesthetized rats

This study analyzed data from 22 simultaneous recordings of single-slice functional magnetic resonance imaging (fMRI; TR = 0.5 s) and bilateral primary somatosensory cortex (S1) local field potentials (LFPs) from 10 male Sprague–Dawley rats (200–300 g; Charles River Laboratories) under dexmedetomidine (DMED) anesthesia, a regime that approximates a lightly sedated or near-awake state (REF). This dataset was originally acquired in multiple previous studies (Pan et al., 2011b) and processed following procedures detailed in (Zhang et al., 2020), which are summarized below.

All experiments were approved by the Emory University Institutional Animal Care and Use Committee and adhered to NIH guidelines. Under DMED anesthesia, fine-tip electrodes (∼10 µm diameter; impedance 1–5 MΩ) were bilaterally implanted in the forelimb regions of S1. Simultaneous single-slice fMRI (9.4 T Bruker MRI system) and bilateral S1 LFP recordings were acquired. Functional imaging targeted a coronal slice covering bilateral S1 regions, with parameters: field of view (FOV) = 1.92 × 1.92 cm², matrix size = 64 × 64, in-plane resolution = 0.3 × 0.3 mm², slice thickness = 2 mm, echo time (TE) = 15 ms, and repetition time (TR) = 500 ms. Each scan lasted 8 min 20 s (1,000 TRs) plus 20 dummy scans for stabilization.

LFP data were low-pass filtered at 100 Hz, downsampled to 500 Hz, and synchronized to fMRI volumes. Gradient artifacts were removed using a template-based subtraction method (Pan et al., 2011b). The cleaned LFPs were band-pass filtered between 0.1 and 100 Hz and divided into six frequency bands: δ (1–4 Hz), θ (4–8 Hz), α (8–12 Hz), low β (12–25 Hz), high β (25–40 Hz), and γ (40–100 Hz). Band-limited power (BLP) time courses were computed using Welch’s method with a 1 s (2 TRs) sliding window and 50% overlap, then normalized for comparability.

fMRI data underwent standard preprocessing, including motion correction and Gaussian spatial smoothing (FWHM = 0.84 mm). Global signals and linear drifts were regressed out, and temporal band-pass filtering (0.01–0.25 Hz) was applied to reduce noise. Subsequently, 22 selected scans were normalized and ROI-based BOLD signals were z-scored. To align with the hemodynamic response function peak latency under DMED (∼2.5 s), BOLD signals were shifted by 2.5 s relative to LFP-derived BLP signals, resulting in 995 TRs (BOLD: [6:1000], BLP: [1:995]) for direct comparison across all frequency bands.

### 4.2. Human motor task fMRI data from the Human Connectome Project (HCP)

Motion task-evoked fMRI (tfMRI) data from the Human Connectome Project (HCP) test-retest cohort were used to assess SWpC sensitivity. The chosen motor task induces robust activations in motor and somatosensory cortices as well as the cerebellum (Barch et al., 2013). Minimally preprocessed 3T motor task fMRI data from the test-retest group was obtained from the Human Connectome Project (HCP). The study included 45 test-retest subjects, each completing two motor task runs. During each run, participants performed five visually cued motions—left foot, right foot, left hand, right hand, and tongue—each repeated twice, resulting in four trials per motion across both runs. Each trial lasted 12 seconds and was preceded by a 3-second visual cue. Additionally, two 12-second resting trials, occurring after one repetition of the five motions, were designated as the null condition for subsequent analysis (Fig. S1).

Task-specific activation maps for the motor task were derived using a three-level FEAT analysis. At *Level 1*, activation maps within each run were identified using a General Linear Model (GLM) as described in (Barch et al., 2013). Five movement predictors—left foot (lf), right foot (rf), left hand (lh), right hand (rh), and tongue (t)—were convolved with a double gamma hemodynamic response function (Glover, 1999), with temporal derivatives included as confounds. The minimally preprocessed fMRI data further underwent spatial smoothing with a FWHM of 4 mm, high-pass filtering with a 200 s cutoff, and prewhitening using FILM to correct for autocorrelations (Woolrich et al., 2001). At *Level 2*, both runs of the Level 1 results for each subject were combined using fixed-effects modeling to produce a single activation map per subject; cluster thresholding was applied with a Z threshold of ±3.29 and a cluster p-threshold of 0.05. At *Level 3*, data across subjects were aggregated using a random-effects model to create group-level activation maps for each task, with cluster thresholding applied using a Z threshold of ±4.42 and a cluster p-threshold of 0.001.

For all three-level analyses, the GLM design included task contrasts for visual cue, each motor task, and each motor task versus baseline. Baseline was defined as the average of all other task conditions (e.g., left hand vs. the average of right hand, left/right foot, and tongue), a method shown to optimize task specificity in brain activation maps (Tripathi et al., 2024). Group-level activation maps for each task contrast relative to its baseline demonstrated consistent task-related activations in the brain. These regions are referred to as task-specific regions of interest (ROIs) throughout this paper. Time series were extracted from these task-specific ROIs for each motion and resting trial, yielding four task time series and four resting time series per subject.

### 4.3. Resting fMRI data in PCVD patients

Post-concussion vestibular dysfunction (PCVD) was hypothesized to involve disrupted directed functional connectivity (FC) along visual–vestibular multisensory processing pathways (Smith et al., 2021). Subacute PCVD (ST) was defined as occurring 2–12 weeks post-concussion and is often considered the stage most responsive to vestibular rehabilitation, whereas chronic PCVD (CH) was defined as vestibular symptoms persisting for >6 months after the concussion. In contrast, healthy controls (HC) had no history of concussion-related vestibular dysfunction or other vestibular impairment. The primary analyses in this study focus on 24 HC versus 24 ST participants. An auxiliary chronic PCVD (CH) cohort (n=24) was included solely to improve the stability of unsupervised brain-state estimation used for state-based feature extraction. All primary hypothesis testing and predictive analyses reported here were restricted to HC vs subacute PCVD (see Section Methods—Functional demonstration on PCVD patients and clinical prediction for details).

Inclusion criteria for PCVD patients required a diagnosis of concussion as defined by the World Health Organization Collaborating Center for Neurotrauma Task Force and clinical evidence of vestibular impairment. Vestibular symptoms were confirmed through subjective reports of dizziness or imbalance, visual motion sensitivity, and symptom provocation during Vestibular/Ocular-Motor Screening (VOMS). Exclusion criteria for both healthy control and PCVD groups included a history of moderate or severe head injury, intracranial hemorrhage, seizure disorder, prior neurologic surgery, peripheral neuropathy, musculoskeletal injuries affecting gait or balance, or chronic drug or alcohol use. Participants with abnormal findings on head impulse testing or videonystagmography (VNG) indicative of peripheral vestibular hypofunction or benign paroxysmal positional vertigo were also excluded.

Resting-state fMRI (rs-fMRI) data were collected for all participants, with each scan lasting 420 seconds and using a repetition time (TR) of 0.7 seconds. Preprocessing was conducted using the CONN Toolbox v19c (Whitfield-Gabrieli and Nieto-Castanon, 2012) following the procedure outlined in (Smith et al., 2021). Standard steps included slice timing, field map correction, motion correction, and registration to the MNI152 template via DARTEL. Motion parameters, outliers, and mean CSF and white matter signals were regressed out. The data were then smoothed with an 8 mm FWHM Gaussian kernel and band-pass filtered (0.01–0.25Hz). All processed data and masks were manually inspected for accuracy, with maximal framewise displacement recorded as 0.76 mm. Timeseries were extracted from 26 parcels of EAGLE449 atlas that cover the core and extended vestibular networks (Smith et al., 2023, Table 1). The extracted timeseries were despiked and low-pass filtered with a cutoff frequency of 0.15 Hz. This dataset provided a foundation for assessing dynamic functional connectivity and spatiotemporal patterns of brain activity, particularly centered around the vestibular networks.

### 4.4. Reliability validation on concurrent LFP-BOLD data in rats

To validate the reliability of SWpC in estimating dynamic directional functional connectivity (FC), we analyzed 22 concurrent LFP-BOLD recordings from the somatosensory cortices (S1L and S1R) of rats (as described in Section 4.1). The physiological expectation of symmetrical information flow between S1L and S1R served as a benchmark for model validation. SWpC was applied to both BOLD signals and band-limited power (BLP) signals across six frequency bands (δ, θ, α, low β, high β, and γ) to evaluate its ability to (1) capture time-varying directional FC at both BOLD and neuronal levels and (2) assess whether the strength and/or duration of information flow detected via BOLD signals was associated with neuronal activity in specific frequency bands. Results were contrasted with metrics derived from traditional sliding window correlation (SWC). Following (Garth John Thompson et al., 2013a), a 50-second sliding window that slids every time point was used for both SWpC and SWC analyses to ensure consistency and facilitate direct comparisons. Each window encompassed 100 timepoints, with a maximum duration of interest set at 15 seconds (or 30 timepoints). To select the optimal model configuration and minimize the risk of overfitting, the Bayesian Information Criterion (BIC) was employed in estimating SWpC durations. This approach enhanced the accuracy and reliability of the duration estimates.

To evaluate SWpC’s performance in estimating time-varying directional FC (S1L→S1R vs. S1R→S1L), we quantified directional asymmetry by calculating the difference in SWpC strength and duration measures between the efferent (S1L→S1R) and afferent (S1R→S1L) directions. For each scan, we first computed the mean SWpC strength or duration across all sliding windows, then defined directional asymmetry as the difference between these two directions. Scan-to-scan variability was determined as the standard deviation of these mean values across all scans for each signal type, including BOLD and BLP signals across six frequency bands (δ, θ, α, low β, high β, and γ). Strength represented the amplitude of directional interactions, while duration reflected how long these interactions persisted within each sliding window. The Wilcoxon signed-rank test was applied to determine whether the median directional asymmetry was significantly lower than the scan-to-scan variability. This non-parametric approach provided a robust assessment of the stability of SWpC-derived estimates, ensuring that results were not disproportionately influenced by deviations from normality or outliers.

We further investigate the neuronal contributions to BOLD-directed FC by correlating SWpC-derived BOLD metrics with BLP signals from six frequency bands. This analysis aimed to clarify whether directional connectivity observed in BOLD signals originated from specific neuronal frequency ranges. To contextualize these findings, we compared SWpC-based correlations with those derived from SWC, highlighting SWpC’s unique capability in characterizing the neural underpinnings of directional information flow. Due to the large variations observed across different scans, a bootstrapping approach was implemented to account for this variability. In each bootstrap iteration, the 22 fMRI scans were randomly divided into two cohorts. For each cohort and each frequency band (δ, θ, α, low β, high β, and γ), Fisher z-transformed correlations were computed between the concatenated sliding window timecourses of BOLD and band-limited power (BLP) signals. This procedure was repeated over 10,000 bootstrap iterations to generate distributions of correlation values for each frequency band. A box plot was employed to visualize the bootstrap distributions, with each box representing the interquartile range (IQR) of the correlations, whiskers extending to 1.5 times the IQR, and outliers displayed as individual points. This approach enabled a robust evaluation of correlation variability while incorporating random variations introduced by cohort splitting.

### 4.5. Sensitivity validation on motor HCP data

To assess the sensitivity of SWpC, we applied it to motor task data from the Human Connectome Project (HCP), which elicits strong directed functional connectivity (FC) between task-specific regions of interest (ROIs). SWpC results were compared with those obtained using standard sliding window correlation (SWC). Each 12-second task or resting trial was treated as an individual window, and both methods were applied to the motion and resting time series for all subjects and runs. Given the small sample size per window, we utilized the corrected Akaike Information Criterion (AICc) for model selection, as it outperforms AIC and BIC under these conditions (Hurvich and Tsai, 1989). For each subject, SWC generated four time-varying strength matrices, and SWpC produced four strength and four duration matrices for each motion-task and corresponding resting condition. Averaging across the four trials of each condition resulted in one time-averaged SWC strength matrix and one SWpC strength and duration matrix per condition.

To assess the asymmetry between directed functional connectivity (FC) results for rest versus task conditions, we quantified asymmetry for each subject’s time-averaged matrix by calculating the Frobenius norm of the difference between the matrix and its transpose, which reflects the overall deviation from symmetry. We conducted normality tests on these asymmetry measures across all subjects to determine the appropriate statistical tests—either independent two-sample t-tests or Mann-Whitney U tests based on data distribution—and calculated Cohen’s d to quantify the magnitude of the observed differences.

Directed FC changes between task and rest conditions were examined for both duration and strength measures. For time-averaged directed FC duration matrices estimated by SWpC, we identified significant differences in directed FC durations between task and rest conditions. Statistical tests were performed on each matrix entry to determine whether these differences were significant (FDR-corrected p-value < 0.01) and exhibited low individual variability (coefficient of variation [CV] < 36%). This approach pinpointed specific connections with significant task-driven changes in the duration of directed FC.

To examine differences in the strength of directed FC between task and rest conditions, we compared time-averaged SWpC results with SWC results by analyzing two pairs of matrices: (directed FC_task, directed FC_rest) and (standard FC_task, standard FC_rest). Element-wise differences for each pair, defined as (directed FC_task - directed FC_rest) and (standard FC_task - standard FC_rest), were computed across all subjects. Statistical approaches were used to quantify these differences. First, significance tests were conducted to identify task-specific connections with low individual variability. For each entry in the difference matrices, a t-test determined whether the mean difference significantly deviated from zero (p < 0.01, FDR-corrected). Among significant connections, those with low individual variability (CV < 30%) were selected. Second, we evaluated the relative sensitivity of SWpC and SWC strengths in detecting task-specific connections. Specifically, the Frobenius norm, defined as the square root of the sum of squared elements in each difference matrix, was calculated to provide a scalar measure of overall distinctions between task and rest. The mean Frobenius norm across all subjects for each matrix pair represented the difference on average. A two-sample t-test was then conducted to determine whether one pair showed significantly larger differences than the other. Additionally, the relative difference between pairs was expressed as the percentage difference in their mean Frobenius norms. This analysis facilitated a direct comparison of SWpC and SWC in detecting task-specific changes in directed FC strength.

Furthermore, we assessed whether time-varying directed FC strength captured trial-to-trial dynamics across the four repetitions of each motion task. We quantified agreement between methods by correlating SWpC- and SWC-derived strength estimates across trials (visualized with a scatter plot). We also examined behavioral relevance by correlating, across the four trials, group-averaged mean directed FC strength (average connectivity across all ROI pairs) with group-averaged reaction time (RT) using Pearson correlation, noting that inference is limited by the small number of time points (n = 4).

### 4.6. Functional demonstration on PCVD patients and clinical prediction

The time-varying causal interactions estimated by SWpC were demonstrated on the rsfMRI data of PCVD patients which are compared with the SWC results. Following previous human study (Allen et al., 2012), we applied a 44-s tapered sliding window that advanced by one TR at each step, generating a time-resolved sequence of SWC and SWpC connectivity matrices for every subject. To obtain a stable, common set of recurring dynamic patterns, the resulting matrices from all available subjects (HC, ST, and CH) were concatenated and submitted to k-means clustering to derive shared brain-state centroids. The Elbow criterion indicated that five clusters provided the most stable decomposition; thus, the cohort was characterized using a common set of five dynamic brain states.

All subsequent inferential and predictive analyses reported in this study were restricted to the primary comparison of HC versus subacute PCVD (ST). After the shared state centroids were estimated, each subject’s windows were assigned to the five states, and we computed (i) connectivity strength, defined as the average edge weight across all ROI pairs within windows assigned to a given state, and (ii) fractional occupation (FO), defined as the proportion of windows belonging to each state. These features captured both the spatial profile and temporal expression of the dynamic patterns and were used for downstream analyses focusing on HC and ST.

To further investigate the large-scale network structure embedded within these dynamic patterns, we averaged the SWC and SWpC matrices across time and applied the Louvain community detection algorithm to the resulting functional networks. Group-level comparisons of modular organization and network segregation/integration were performed for HC versus ST only, in order to assess whether SWpC and SWC highlighted different vestibular–multisensory pathway organization and whether subacute PCVD showed altered network topology relative to controls.

Finally, to evaluate the clinical relevance of state-derived features given the limited patient dataset, we implemented a nested leave-one-out cross-validation (LOOCV) machine-learning framework (Vergara et al., 2018) to discriminate subacute PCVD (ST) from HC. In each fold, one participant was held out as the test case, and a support vector machine (SVM) classifier was trained on the remaining participants using the features derived from the five states for each of the three cases: SWC strength, SWpC strength, and SWpC duration. Within each training fold, features were ranked by their two-sample t-test statistic between groups, and the top [2%: 2%: 20%] were retained. The optimal threshold was selected within the inner loop of the nested CV procedure. The resulting classifier was evaluated on the held-out participant, and performance was quantified using the area under the ROC curve (AUC). This nested design prevented information leakage between training and testing and provided an unbiased estimate of the ability of dynamic connectivity features to discriminate subacute PCVD patients from healthy controls.

## 5. Data and code availability statement

Our code for computing SWpC is available at https://github.com/inspirelab-site/swpc.

## 6. Credit authorship contribution statement

**Nan Xu:** Conceptualization, Data curation, Data preprocessing, Methodology, Formal analysis, Writing – original draft, Writing – review & editing, Funding acquisition. **Xiaodi Zhang:** Data curation, Data preprocessing for rodent data, **Wen-Ju Pan:** Rodent data acquisition. **Jeremy L. Smith:** Data curation, Data preprocessing for PCVD data. **Eric H. Schumacher:** Writing – review & editing. **Jason W. Allen:** Writing – review & editing, Supervision. **Vince D. Calhoun:** Writing – review & editing, Supervision. **Shella D. Keilholz:** Writing – review & editing, Supervision.

## 8. Acknowledgement

Nan Xu thanks the funding support for National Institutes of Health (NIH K99/R00NS123113).

## References

Aedo-Jury, F., Schwalm, M., Hamzehpour, L., Stroh, A., 2020. Brain states govern the spatio-temporal dynamics of resting state functional connectivity. Elife 9. 10.7554/eLife.53186

Allen, E.A., Damaraju, E., Plis, S.M., Erhardt, E.B., Eichele, T., Calhoun, V.D., 2012. Tracking whole-brain connectivity dynamics in the resting state. Cereb Cortex 24, 663–676. 10.1093/cercor/bhs352

Barch, D.M., Burgess, G.C., Harms, M.P., Petersen, S.E., Schlaggar, B.L., Corbetta, M., Glasser, M.F., Curtiss, S., Dixit, S., Feldt, C., Nolan, D., Bryant, E., Hartley, T., Footer, O., Bjork, J.M., Poldrack, R., Smith, S., Johansen-Berg, H., Snyder, A.Z., Van Essen, D.C., 2013. Function in the human connectome: task-fMRI and individual differences in behavior. Neuroimage 80, 169–189. 10.1016/J.NEUROIMAGE.2013.05.033

Chaudhuri, R., Knoblauch, K., Gariel, M.A., Kennedy, H., Wang, X.J., 2015. A Large-Scale Circuit Mechanism for Hierarchical Dynamical Processing in the Primate Cortex. Neuron 88, 419–431. 10.1016/J.NEURON.2015.09.008/ATTACHMENT/277AD0E9-7600-4E84-B278-784BAFBC3B1D/MMC3.PDF

Fieblinger, T., 2021. Striatal Control of Movement: A Role for New Neuronal (Sub-) Populations? Front. Hum. Neurosci. 15, 697284. 10.3389/FNHUM.2021.697284/BIBTEX

Frank, S.M., Greenlee, M.W., 2018. The parieto-insular vestibular cortex in humans: More than a single area? J. Neurophysiol. 120, 1438–1450. 10.1152/JN.00907.2017/ASSET/IMAGES/LARGE/Z9K0091847620004.JPEG

Fries, P., 2015. Rhythms for Cognition: Communication through Coherence. Neuron 88, 220–235. 10.1016/J.NEURON.2015.09.034/ASSET/2E400083-7871-4FD3-8A27-D8163B6B7441/MAIN.ASSETS/GR9.JPG

Friston, K.J., 2011. Functional and Effective Connectivity: A Review. Brain Connect. 1, 13–36. 10.1089/brain.2011.0008

Friston, K.J., Harrison, L., Penny, W., 2003. Dynamic causal modelling. Neuroimage 19, 1273–1302. 10.1016/S1053-8119(03)00202-7

Glover, G.H., 1999. Deconvolution of Impulse Response in Event-Related BOLD fMRI1. Neuroimage 9, 416–429. 10.1006/NIMG.1998.0419

Handwerker, D.A., Ollinger, J.M., D’Esposito, M., 2004. Variation of BOLD hemodynamic responses across subjects and brain regions and their effects on statistical analyses. Neuroimage 21, 1639–1651. 10.1016/J.NEUROIMAGE.2003.11.029

Hurvich, C.M., Tsai, C.L., 1989. Regression and time series model selection in small samples. Biometrika 76, 297–307. 10.1093/BIOMET/76.2.297

Hutchison, R M, Womelsdorf, T., Allen, E.A., Bandettini, P.A., Calhoun, V.D., Corbetta, M., Della Penna, S., Duyn, J.H., Glover, G.H., Gonzalez-Castillo, J., Handwerker, D.A., Keilholz, S., Kiviniemi, V., Leopold, D.A., de Pasquale, F., Sporns, O., Walter, M., Chang, C., 2013. Dynamic functional connectivity: promise, issues, and interpretations. Neuroimage 80, 360–378. 10.1016/j.neuroimage.2013.05.079

Hutchison, R. Matthew, Womelsdorf, T., Allen, E.A., Bandettini, P.A., Calhoun, V.D., Corbetta, M., Della Penna, S., Duyn, J.H., Glover, G.H., Gonzalez-Castillo, J., Handwerker, D.A., Keilholz, S., Kiviniemi, V., Leopold, D.A., de Pasquale, F., Sporns, O., Walter, M., Chang, C., 2013. Dynamic functional connectivity: Promise, issues, and interpretations. Neuroimage 80, 360–378. 10.1016/J.NEUROIMAGE.2013.05.079

Indovina, I., Bosco, G., Riccelli, R., Maffei, V., Lacquaniti, F., Passamonti, L., Toschi, N., 2020. Structural connectome and connectivity lateralization of the multimodal vestibular cortical network. Neuroimage 222, 117247. 10.1016/J.NEUROIMAGE.2020.117247

Keilholz, S.D., Magnuson, M.E., Pan, W.J., Willis, M., Thompson, G.J., 2013. Dynamic properties of functional connectivity in the rodent. Brain Connect 3, 31–40. 10.1089/brain.2012.0115

Liang, Z., Liu, X., Zhang, N., 2015. Dynamic resting state functional connectivity in awake and anesthetized rodents. Neuroimage 104, 89–99. 10.1016/j.neuroimage.2014.10.013

Manto, M., Bower, J.M., Conforto, A.B., Delgado-García, J.M., Da Guarda, S.N.F., Gerwig, M., Habas, C., Hagura, N., Ivry, R.B., Marien, P., Molinari, M., Naito, E., Nowak, D.A., Ben Taib, N.O., Pelisson, D., Tesche, C.D., Tilikete, C., Timmann, D., 2011. Consensus Paper: Roles of the Cerebellum in Motor Control—The Diversity of Ideas on Cerebellar Involvement in Movement. The Cerebellum 2011 11:2 11, 457–487. 10.1007/S12311-011-0331-9

Medrano, J., Friston, K.J., Zeidman, P., 2024. Dynamic Causal Models of Time-Varying Connectivity. arXiv:2411.16582.

Mitra, A, Snyder, A.Z., Blazey, T., Raichle, M.E., 2015. Lag threads organize the brain’s intrinsic activity. Proc Natl Acad Sci U S A 112, E2235–44. 10.1073/pnas.1503960112

Mitra, Anish, Snyder, A.Z., Blazey, T., Raichle, M.E., 2015. Lag threads organize the brain’s intrinsic activity. Proc. Natl. Acad. Sci. U. S. A. 112, E2235–E2244. 10.1073/PNAS.1503960112/SUPPL_FILE/PNAS.1503960112.SM04.MP4

Moon, H.S., Vo, T.T., Im, G.H., Hong, S.J., Kim, S.G., 2025. Interhemispheric resting-state functional connectivity correlates with spontaneous neural interactions. Proc. Natl. Acad. Sci. U. S. A. 122, e2505294122. 10.1073/PNAS.2505294122/SUPPL_FILE/PNAS.2505294122.SAPP.PDF

Murray, J.D., Bernacchia, A., Freedman, D.J., Romo, R., Wallis, J.D., Cai, X., Padoa-Schioppa, C., Pasternak, T., Seo, H., Lee, D., Wang, X.J., 2014. A hierarchy of intrinsic timescales across primate cortex. Nature Neuroscience 2014 17:12 17, 1661–1663. 10.1038/nn.3862

Pan, W. ju, Thompson, G., Magnuson, M., Majeed, W., Jaeger, D., Keilholz, S., 2011. Broadband local field potentials correlate with spontaneous fluctuations in functional magnetic resonance imaging signals in the rat somatosensory cortex under isoflurane Anesthesia. Brain Connect. 1, 119–131. 10.1089/brain.2011.0014

Park, H.J., Friston, K.J., Pae, C., Park, B., Razi, A., 2018. Dynamic effective connectivity in resting state fMRI. Neuroimage 180, 594–608. 10.1016/j.neuroimage.2017.11.033

Park, J., Xu, N., Nezafati, M., 2026. A comprehensive analysis of brain network complexity in task-based fMRI using entropy: systematic review. Brain Imaging and Behavior 2026 20:2 20, 61-. 10.1007/S11682-026-01124-Y

Peters, J., Janzing, D., Scholkopf, B., 2017. Elements of Causal Inference: Foundations and Learning Algorithms. MIT press.

Rizzolatti, G., Luppino, G., 2001. The cortical motor system. Neuron 31, 889–901. 10.1016/S0896-6273(01)00423-8

Sasegbon, A., Hamdy, S., 2023. The Role of the Cerebellum in Swallowing. Dysphagia 38, 497–509. 10.1007/S00455-021-10271-X/FIGURES/5

Seth, A.K., Barrett, A.B., Barnett, L., 2015. Granger Causality Analysis in Neuroscience and Neuroimaging. Journal of Neuroscience 35, 3293–3297. 10.1523/JNEUROSCI.4399-14.2015

Shakil, S., Lee, C.H., Keilholz, S.D., 2016. Evaluation of sliding window correlation performance for characterizing dynamic functional connectivity and brain states. Neuroimage 133, 111–128. 10.1016/j.neuroimage.2016.02.074

Smith, J.L., Ahluwalia, V., Gore, R.K., Allen, J.W., 2023. Eagle-449: A volumetric, whole-brain compilation of brain atlases for vestibular functional MRI research. Scientific Data 2023 10:1 10, 1–13. 10.1038/s41597-023-01938-1

Smith, J.L., Trofimova, A., Ahluwalia, V., Casado Garrido, J.J., Hurtado, J., Frank, R., Hodge, A., Gore, R.K., Allen, J.W., 2021. The “vestibular neuromatrix”: A proposed, expanded vestibular network from graph theory in post-concussive vestibular dysfunction. Hum. Brain Mapp. 10.1002/HBM.25737

Smith, Stephen M., Miller, K.L., Salimi-Khorshidi, G., Webster, M., Beckmann, C.F., Nichols, T.E., Ramsey, J.D., Woolrich, M.W., 2011. Network modelling methods for FMRI. Neuroimage 54, 875–891. 10.1016/J.NEUROIMAGE.2010.08.063

Smith, S M, Miller, K.L., Salimi-Khorshidi, G., Webster, M., Beckmann, C.F., Nichols, T.E., Ramsey, J.D., Woolrich, M.W., 2011. Network modelling methods for FMRI. Neuroimage 54, 875–891. 10.1016/j.neuroimage.2010.08.063

Thompson, G.J., Merritt, M.D., Pan, W.-J., Magnuson, M.E., Grooms, J.K., Jaeger, D., Keilholz, S.D., 2013. Neural correlates of time-varying functional connectivity in the rat. Neuroimage 83, 826–836. 10.1016/j.neuroimage.2013.07.036

Todorov, E., 2004. Optimality principles in sensorimotor control. Nature Neuroscience 2004 7:9 7, 907–915. 10.1038/nn1309

Tripathi, V., Rigolo, L., Bracken, B.K., Galvin, C.P., Golby, A.J., Tie, Y., Somers, D.C., 2024. Utilizing connectome fingerprinting functional MRI models for motor activity prediction in presurgical planning: A feasibility study. Hum. Brain Mapp. 45. 10.1002/HBM.26764

van den Heuvel, M.P., Hulshoff Pol, H.E., 2010. Exploring the brain network: A review on resting-state fMRI functional connectivity. European Neuropsychopharmacology 20, 519–534. 10.1016/J.EURONEURO.2010.03.008

Vergara, V.M., Mayer, A.R., Kiehl, K.A., Calhoun, V.D., 2018. Dynamic functional network connectivity discriminates mild traumatic brain injury through machine learning. Neuroimage Clin. 19, 30–37. 10.1016/J.NICL.2018.03.017

Whitfield-Gabrieli, S., Nieto-Castanon, A., 2012. Conn: A Functional Connectivity Toolbox for Correlated and Anticorrelated Brain Networks. https://home.liebertpub.com/brain 2, 125–141. 10.1089/BRAIN.2012.0073

Woolrich, M.W., Ripley, B.D., Brady, M., Smith, S.M., 2001. Temporal autocorrelation in univariate linear modeling of FMRI data. Neuroimage 14, 1370–1386. 10.1006/NIMG.2001.0931

Xu, N., Doerschuk, P.C., Keilholz, S.D., Spreng, R.N., 2021a. Spatiotemporal functional interactivity among large-scale brain networks. Neuroimage 227, 117628. 10.1016/J.NEUROIMAGE.2020.117628

Xu, N., Doerschuk, P.C., Keilholz, S.D., Spreng, R.N., 2021b. Spatiotemporal functional interactivity among large-scale brain networks. Neuroimage 227, 117628. 10.1016/j.neuroimage.2020.117628

Xu, N., Spreng, R.N., Doerschuk, P.C., 2017. Initial validation for the estimation of resting-state fMRI effective connectivity by a generalization of the correlation approach. Front. Neurosci. 11, 271. 10.3389/fnins.2017.00271

Zhang, X., Pan, W.J., Keilholz, S.D., 2020. The relationship between BOLD and neural activity arises from temporally sparse events. Neuroimage 207, 116390. 10.1016/j.neuroimage.2019.116390

