## Supplementary Materials for "Time-resolved directional organization of brain networks: distinct strength and temporal signatures from neurophysiology to disease"

Supplementary Material

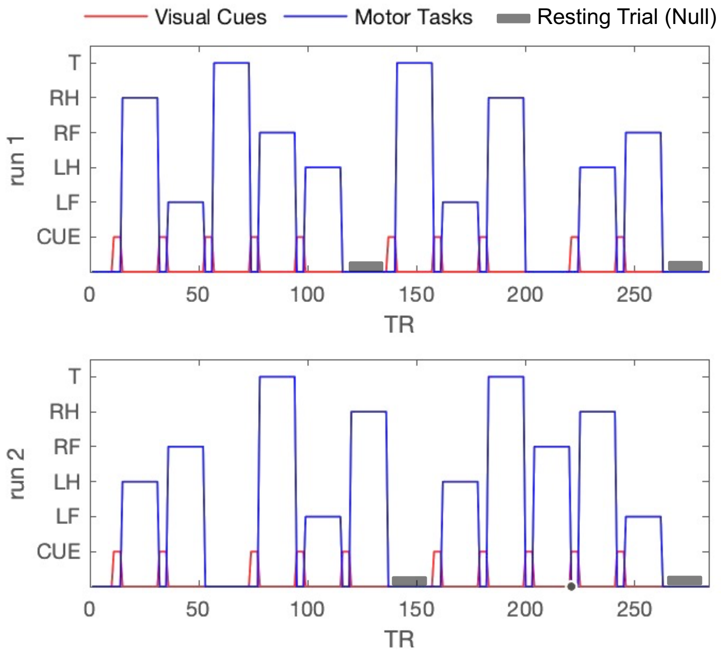


Fig. S1. Timing of the visually cued motion tasks and corresponding resting trials (null conditions) for each fMRI run.

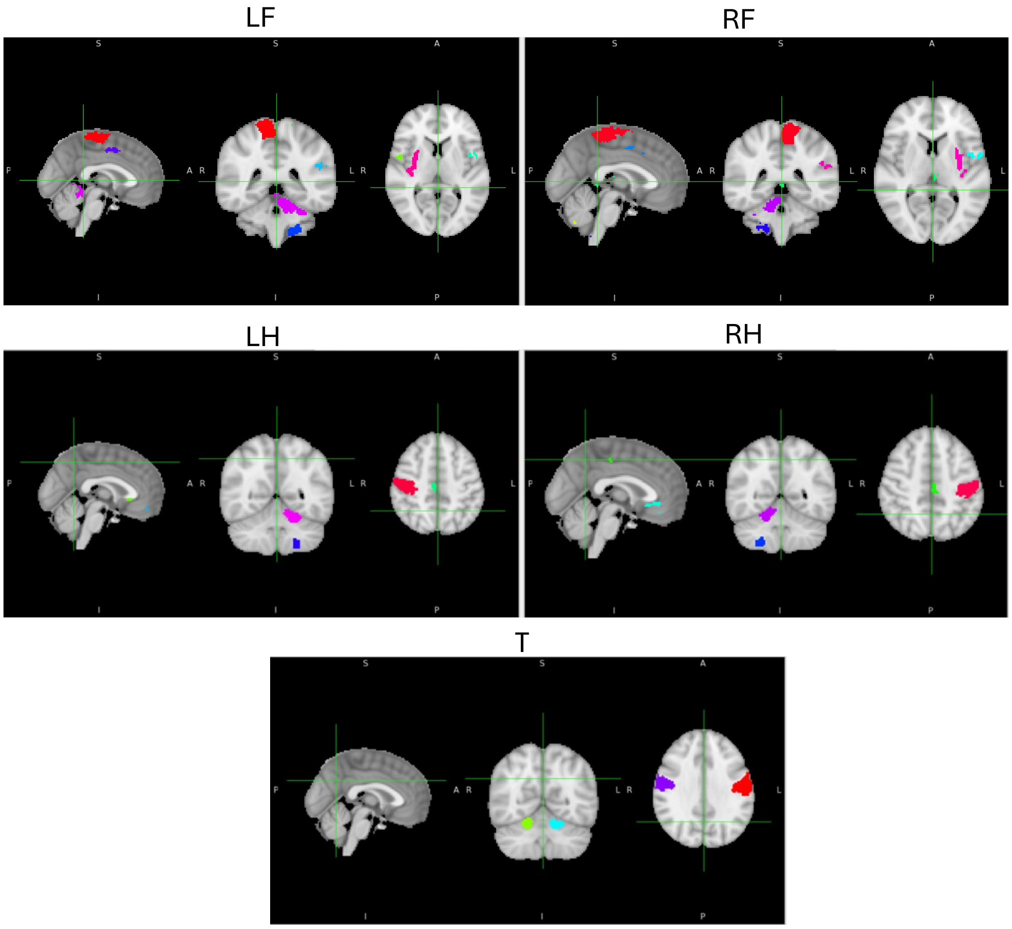


Fig. S2. Activation maps (ROIs) derived from FEAT analysis for each motion task.

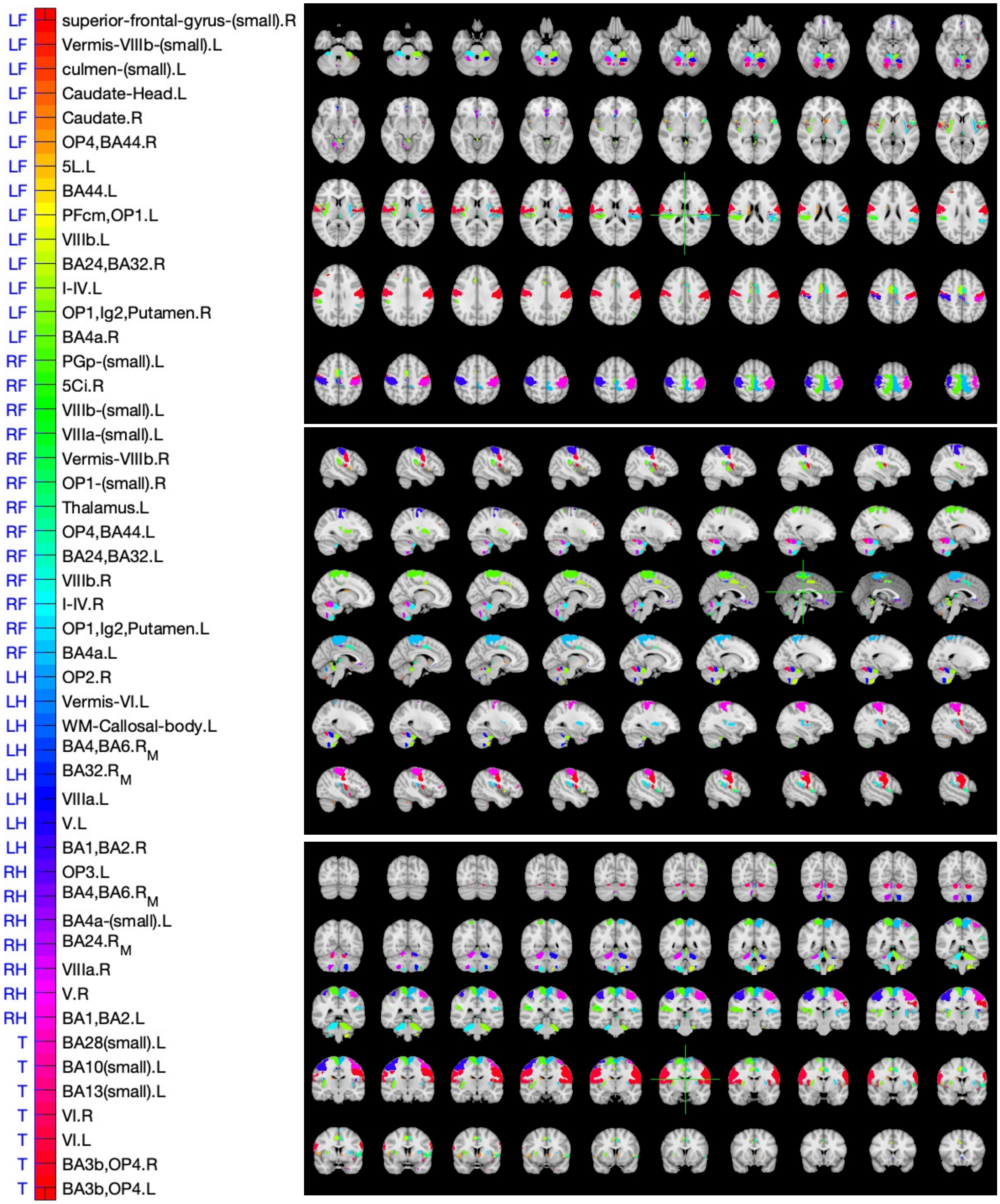


Fig. S3. 49 motion-specific ROIs identified by 3-level FEAT analysis

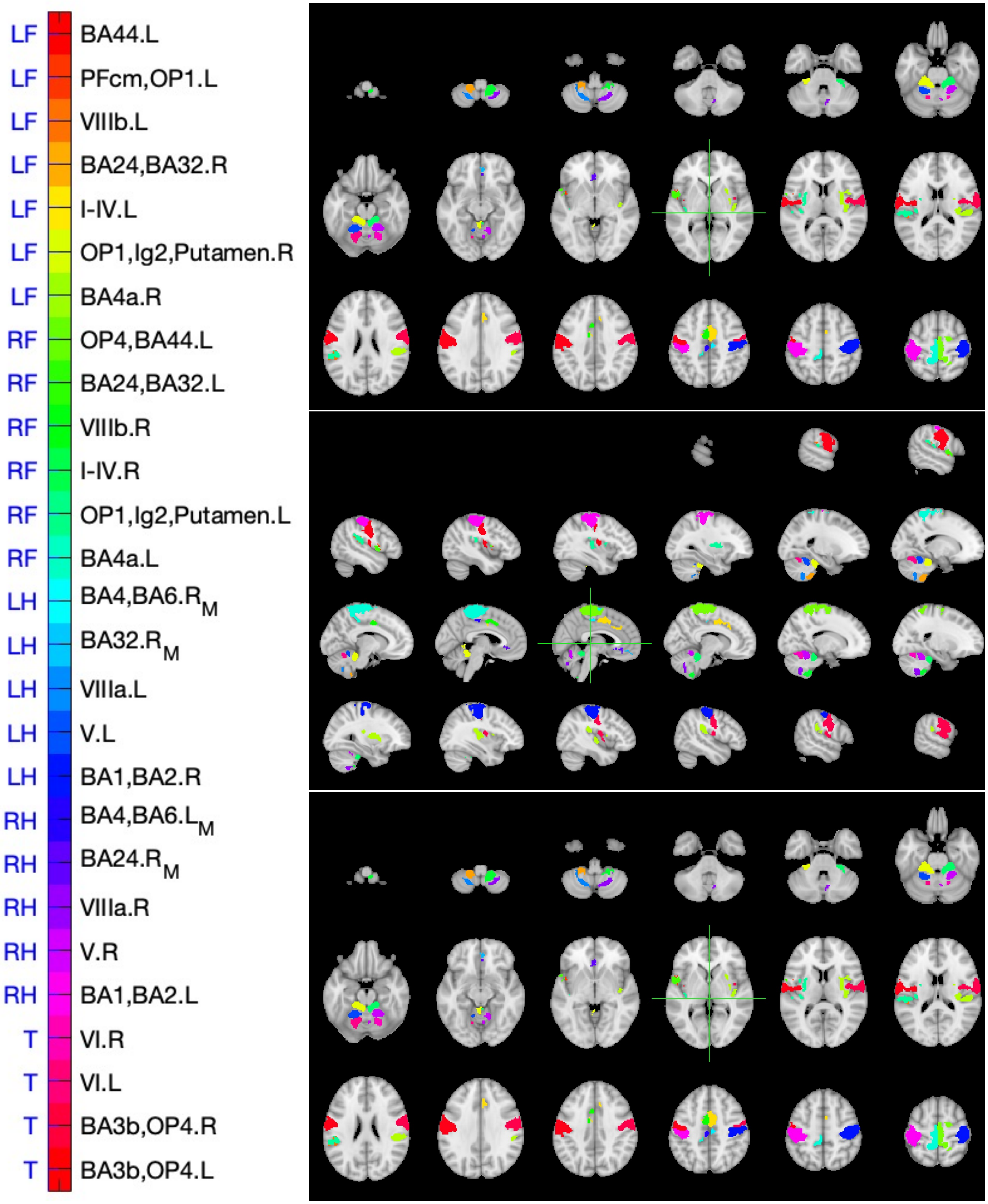


Fig. S4. 27 selected large motion-specific ROIs for illustration purposes. These include 13 foot ROIs, 10 hand ROIs, and 4 tongue ROIs, with smaller ROIs excluded (fewer than 100 voxels for foot motions and fewer than 30 voxels for hand and tongue motions). These 27 ROIs exhibit approximate symmetry for left/right foot and hand tasks.

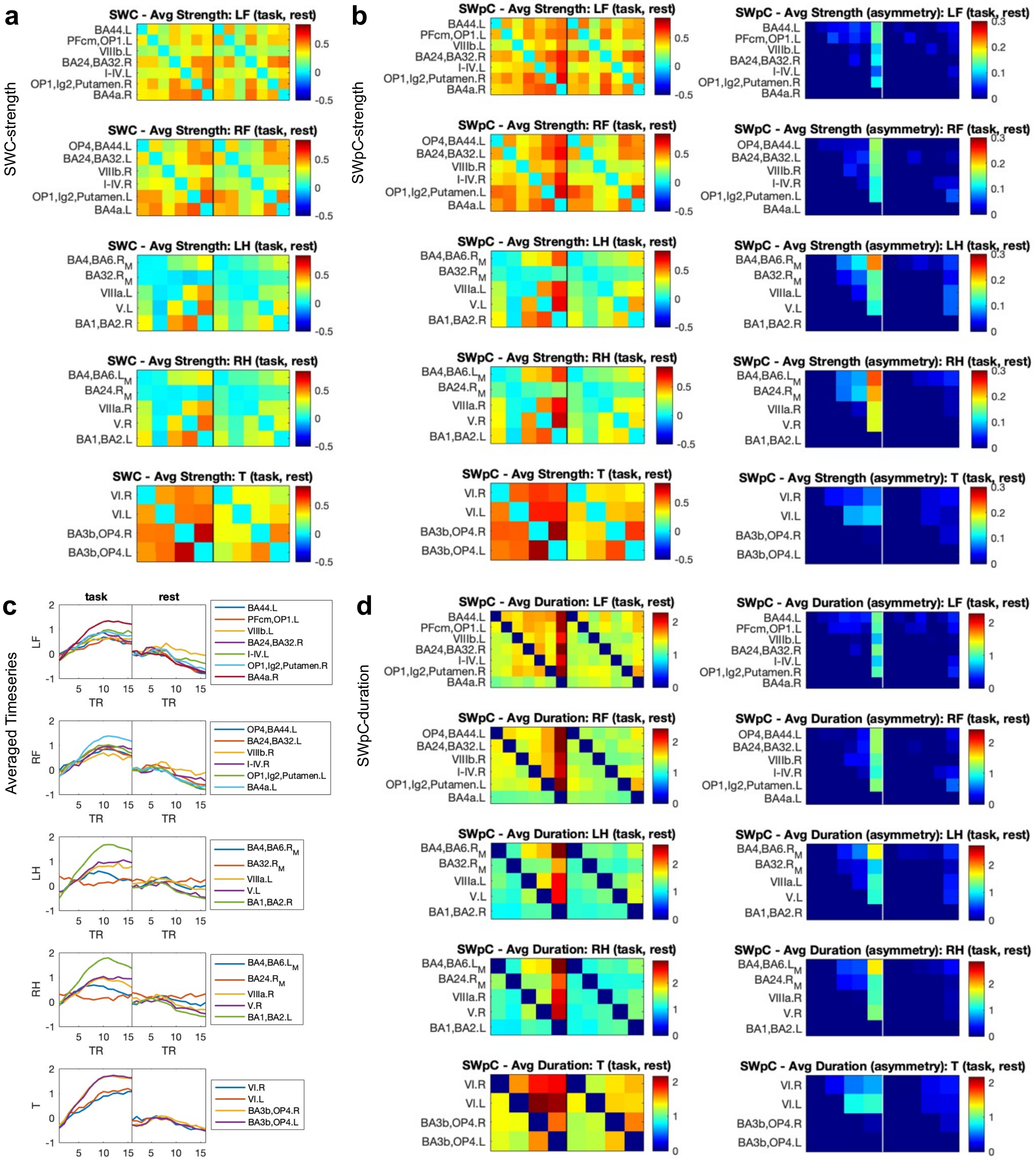


Fig. S5. Group-averaged (directed) FC strength and duration matrices as well as fMRI timeseries during task and rest trials.

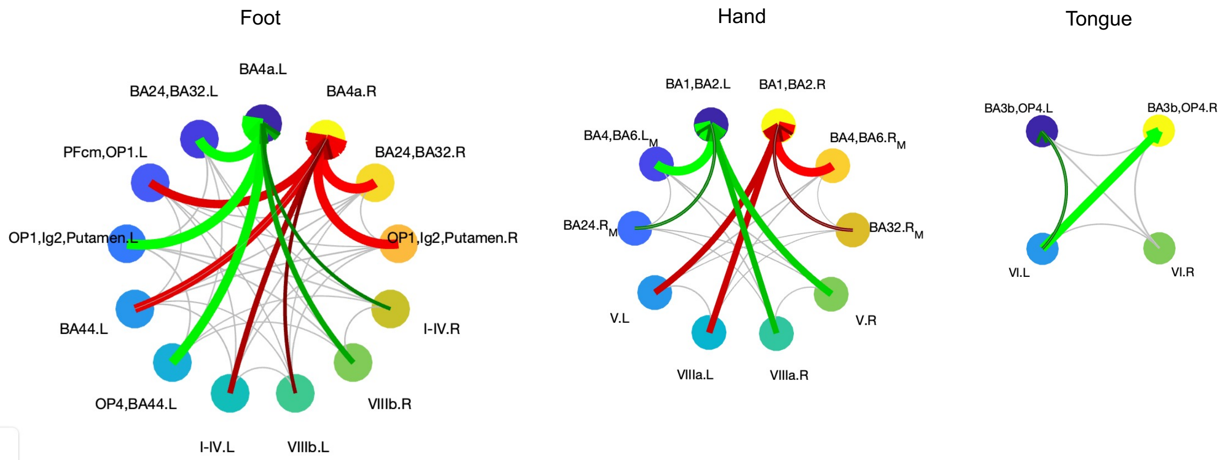


Fig. S6. Connectograms of directed FC with significant differences in duration between task and rest for hand (left), (middle), and tongue (right) motions. Red (green) arrows indicate task vs. rest differences for left (right) motion with low individual variability (CV < 36%). For tongue motions, green arrows represent significant evoked directed FC strengths with low variability. Gray connections denote all predicted directed connections from SWpC. To enhance clarity, connectograms include 13 foot ROIs, 10 hand ROIs, and 4 tongue ROIs, excluding smaller ROIs (fewer than 100 voxels for foot motions and fewer than 30 voxels for hand and tongue motions). The remaining ROIs exhibit approximate symmetry for left/right foot and hand tasks. Under these thresholds, tongue-evoked information flow among the 4 larger ROIs does not form hubs in the motor cortex but connects VI.L to two somatomotor cortical regions, potentially due to the applied threshold amount the limited number of connections.

| | Motion | Distribution Type | Asymmetry Measure (Task) | Asymmetry Measure (Rest) | P-value | Cohen's D effect size |
|---|---|---|---|---|---|---|
| **strength** | LF | Normal | 1.87 | 1.50 | 1.53E-06 | 1.0712 |
| | RF | Normal | 1.66 | 1.39 | 1.87E-06 | 1.0499 |
| | LH | Normal | 1.13 | 0.87 | 3.23E-06 | 1.0223 |
| | RH | Normal | 1.20 | 0.71 | 2.43E-12 | 1.7892 |
| | T | Normal | 0.99 | 0.68 | 8.03E-07 | 1.1007 |
| **duration** | LF | Normal | 10.21 | 7.31 | 8.19E-08 | 1.246 |
| | RF | Normal | 9.45 | 6.49 | 3.06E-13 | 1.8447 |
| | LH | Normal | 6.66 | 3.91 | 8.99E-13 | 1.8305 |
| | RH | Normal | 7.33 | 3.14 | 3.15E-15 | 2.1781 |
| | T | Normal | 6.67 | 3.32 | 1.37E-13 | 1.9559 |

Table S1. Directional Asymmetry Measures in SWpC strength and Durations.

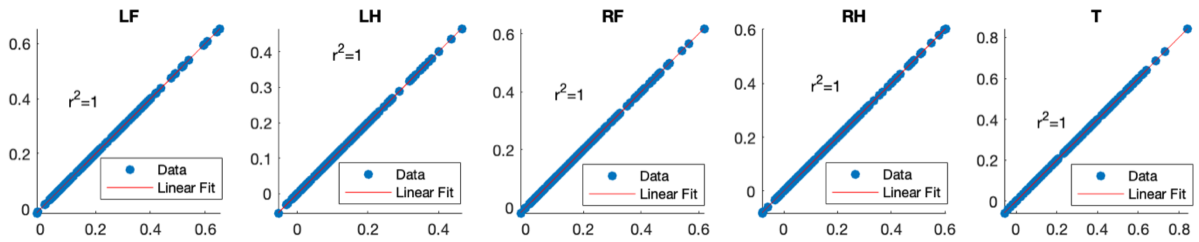


Fig S7. Scatter Plot of SWPC-Directed FC Matrix Strength vs. SWC FC Matrix Strength

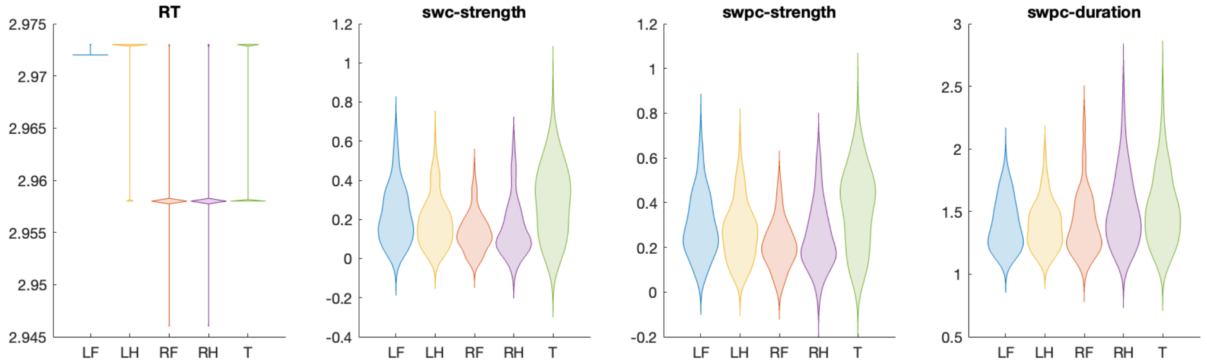


Fig S8. Violin plots of reaction time (RT), and SWC as well as SWpC estimates.

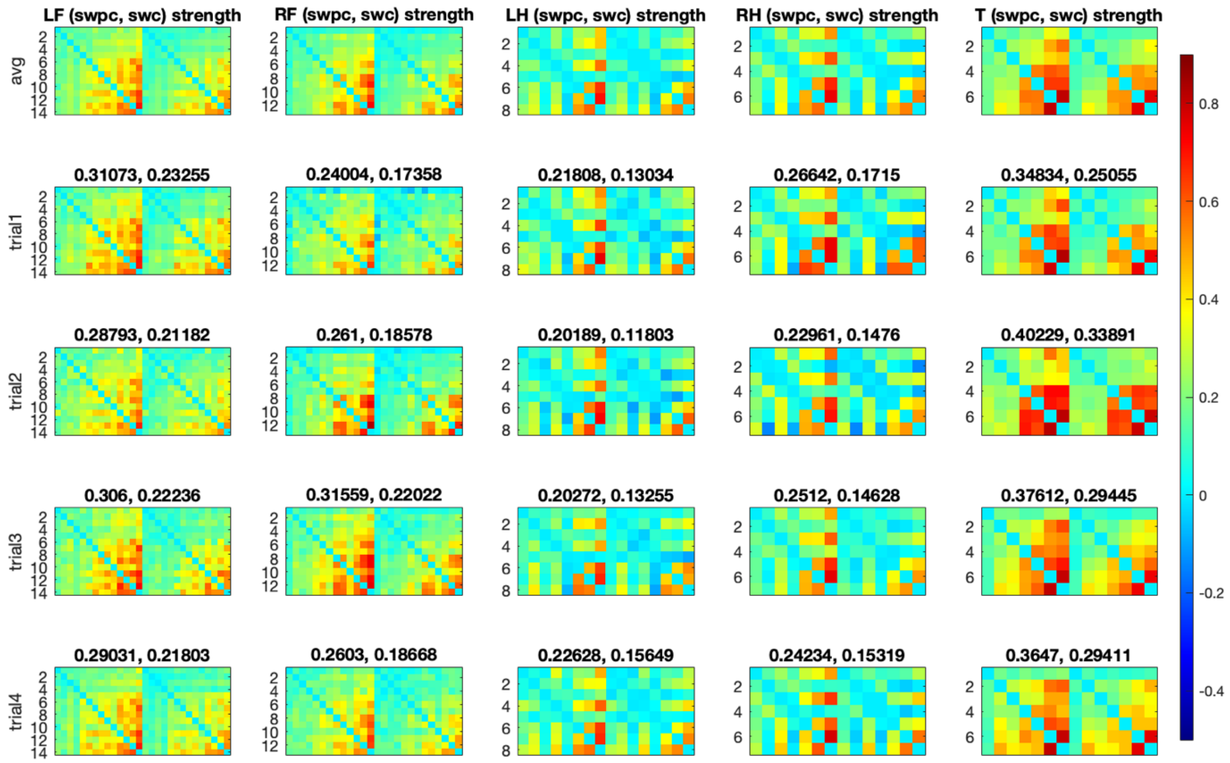


Fig S9. Group-averaged SWpC and SWC strength during each task trial. The mean FC strength was listed on the top of each matrix.

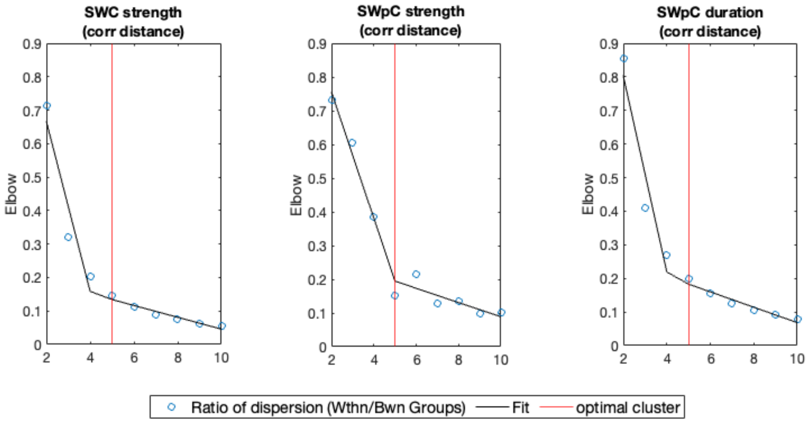


Fig S10. Optimal k-means cluster number by Elbow method.

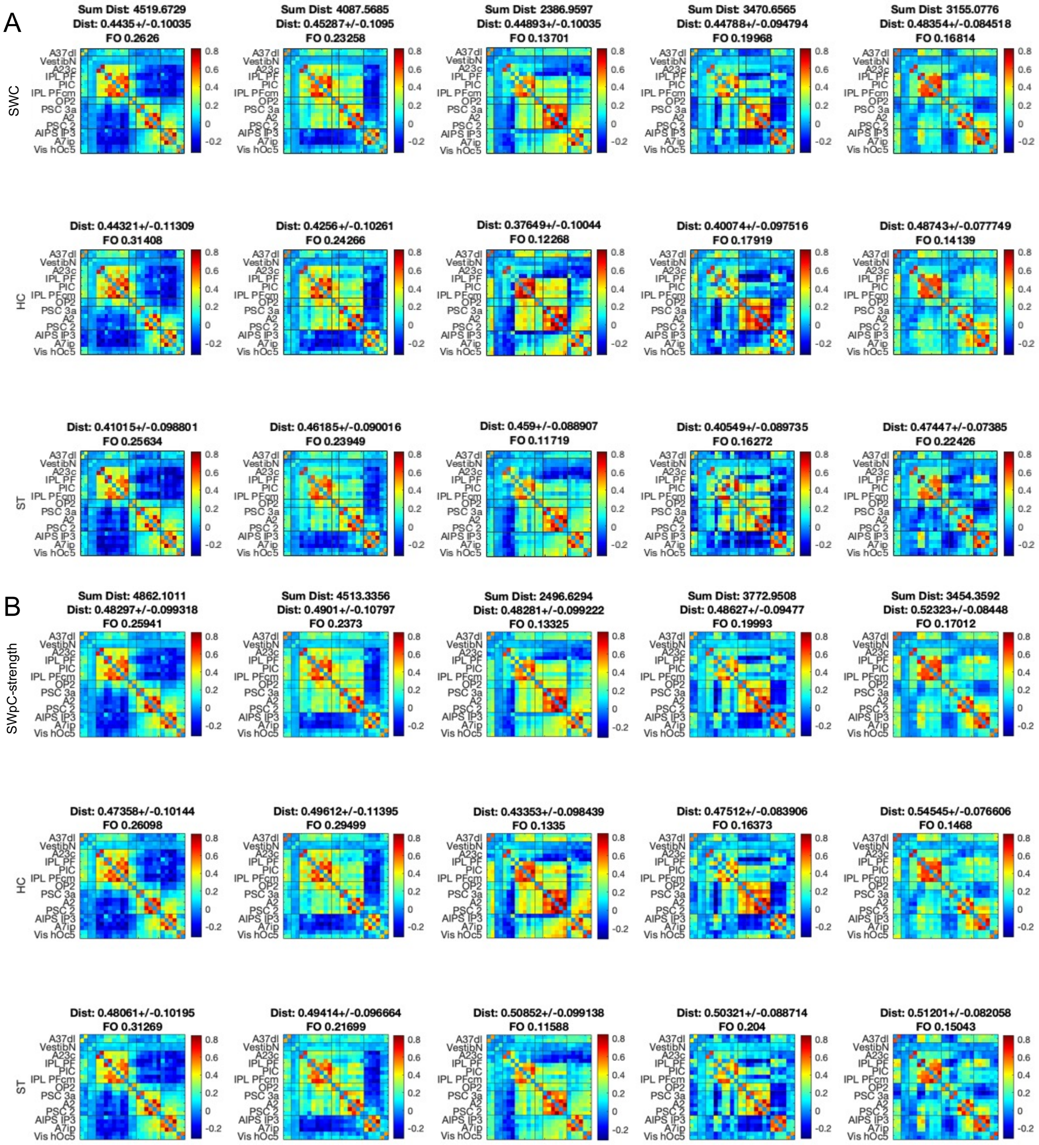

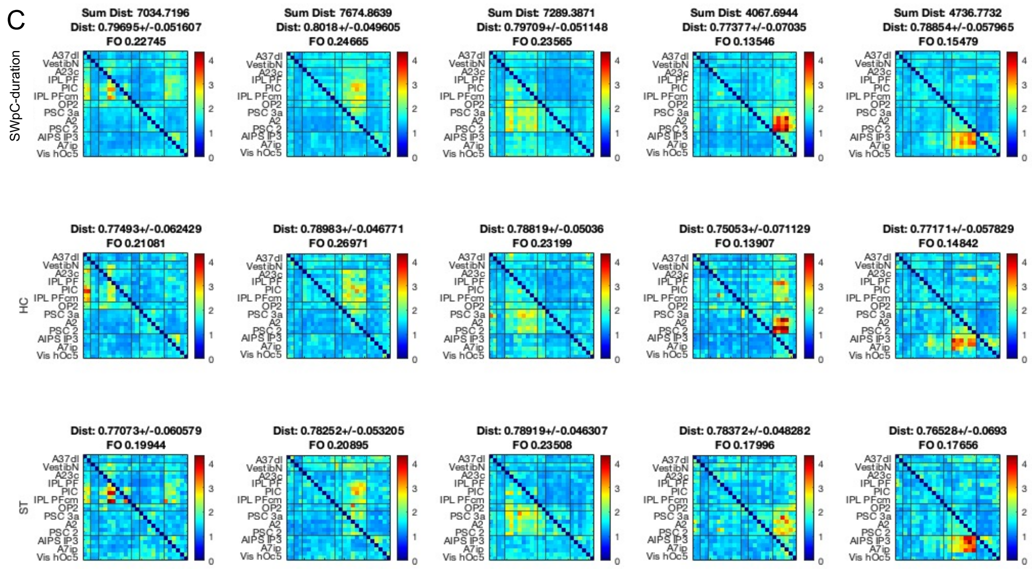


Fig. S12. Brain states estimated using (A) SWC, (B) SWpC-strength, and (C) SWpC-duration, shown for states shared across HC, ST, and CH (top), as well as states specific to HC (middle) and ST (bottom).

| | State 1 | State 2 | State 3 | State 4 | State 5 |
|---|---|---|---|---|---|
| HC | 27% (23) | 29% (24) | 14% (17) | 16% (16) | 14% (21) |
| | 26% (23) | 30% (23) | 13% (20) | 16% (17) | 15% (20) |
| | 22% (24) | 30% (24) | 22% (24) | 13% (24) | 14% (24) |
| ST | 31% (23) | 21% (22) | 11% (19) | 21% (22) | 15% (20) |
| | 31% (23) | 22% (22) | 12% (19) | 20% (21) | 15% (20) |
| | 23% (24) | 21% (24) | 24% (24) | 13% (24) | 19% (24) |

Table S2 Fraction of occupation for each state for each group with parathesis including the number of subjects that visit each state in each group. Color-code (black, blue and purple) enlists the results of swc-strength, swpc-strength, swpc-duration.

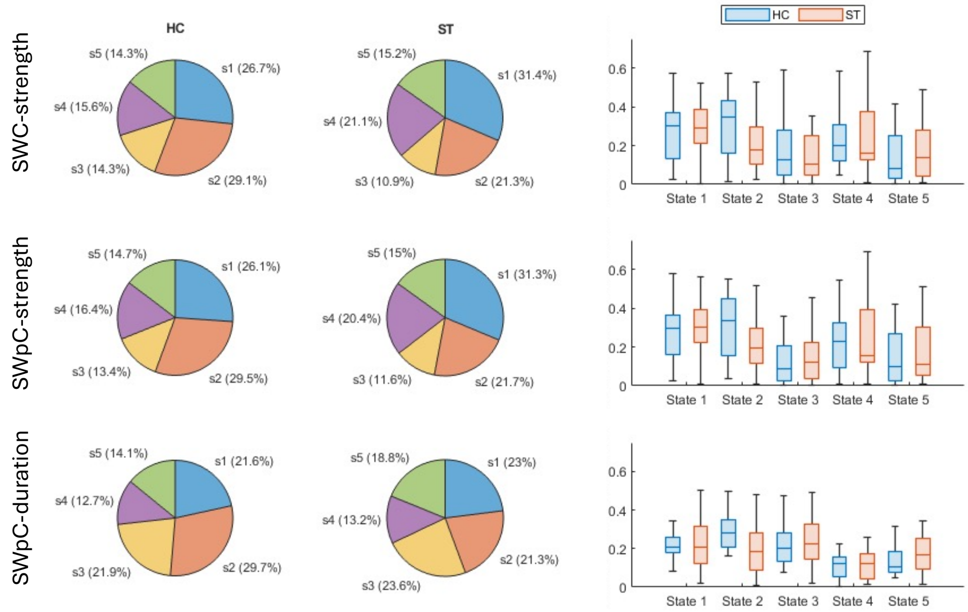


Fig. S13. Fractional occupancy in Healthy Controls and subacute PCVD patients. Left: pie charts based on group mean values. Right: bar plots showing the group means and variability.

| p-values | state 1 | state 2 | state 3 | state 4 | state 5 |
|---|---|---|---|---|---|
| SWC-strength | 0.376 | 0.121 | 0.496 | 0.294 | 0.864 |
| SWpC-strength | 0.330 | 0.131 | 0.711 | 0.444 | 0.946 |
| SWpC-duration | 0.673 | **0.035** | 0.614 | 0.857 | 0.152 |

Table S3. Two-sample t-test p-values comparing groups for each state (States 1–5) across SWC strength, SWpC strength, and SWpC duration.
